# Ductal pancreatic cancer interception by FGFR2 abrogation

**DOI:** 10.1101/2024.10.16.618726

**Authors:** Claudia Tonelli, Astrid Deschênes, Victoria Gaeth, Amanda Jensen, Nandan Vithlani, Melissa A. Yao, Zhen Zhao, Youngkyu Park, David A. Tuveson

## Abstract

Activating KRAS mutations are a key feature of pancreatic ductal adenocarcinoma (PDA) and drive tumor initiation and progression. However, mutant KRAS by itself is weakly oncogenic. The pathways that cooperate with mutant KRAS to induce tumorigenesis are less-defined. Analyzing organoids and murine and human pancreatic specimens, we found that the receptor tyrosine kinase FGFR2 was progressively up-regulated in mutant KRAS-driven metaplasia, pre-neoplasia and Classical PDA. Using genetic mouse models, we showed that FGFR2 supported mutant KRAS-driven transformation of acinar cells by promoting proliferation and MAPK pathway activation. FGFR2 abrogation significantly delayed tumor formation and extended the survival of these mice. Furthermore, we discovered that FGFR2 collaborated with EGFR and dual blockade of these receptor signaling pathways significantly reduced mutant KRAS-induced pre-neoplastic lesion formation.

Together, our data have uncovered a pivotal role for FGFR2 in the early phases of pancreatic tumorigenesis, paving the way for future therapeutic applications of FGFR2 inhibitors for pancreatic cancer interception.

**STATEMENT OF SIGNIFICANCE:** Mutant KRAS-expressing pancreatic intraepithelial neoplasias (PanINs), the precursor lesions of PDA, are prevalent in the average healthy adult but rarely advance to invasive carcinoma. Here, we discovered that FGFR2 promoted PDA progression by amplifying mutant KRAS signaling and that inactivation of FGFR2 intercepted disease progression.

## INTRODUCTION

Pancreatic ductal adenocarcinoma (PDA) remains one of the deadliest malignancies with a 5 year survival rate of only 13% (1). Activating mutations of *KRAS* in the exocrine pancreas constitute the prevalent initiating event and drive the formation of precursor lesions called pancreatic intraepithelial neoplasia (PanIN) (2). Analysis of pancreata from organ donors revealed that PanINs are common in the general healthy population and unlikely to progress to infiltrating carcinoma, given the relatively low incidence of PDA (3). Additional signaling pathways, among which epidermal growth factor receptor (EGFR) is the most well studied (4,5), cooperate with mutant KRAS to drive PDA initiation and progression. Inactivating mutations of tumor suppressor genes (eg, *TP53*, *CDKN2A*, *SMAD4*) ultimately drive tumor formation (6,7).

The extended time required for the progression of PanIN to PDA provides a window of opportunity to intervene and block cancer development. Understanding the mechanisms that promote the early phases of pancreatic tumorigenesis is key to enable the development of strategies for better pancreatic cancer risk assessment, prevention and interception (8).

The fibroblast growth factor receptor (FGFR) gene family consists of four members, FGFR1 to FGFR4 (9). FGFRs are tyrosine kinase receptors (RTKs) whose downstream pathways include the RAS/RAF-MAPK, PI3K/AKT, PLCγ and STAT. Context-dependent expression, ligand binding capacities and alternative splicing isoforms of the FGFRs determine differences in the pathways induced and consequent phenotypic changes. FGFRs are aberrantly activated through mutations, gene fusions and copy number amplifications in 5–10% of all human cancers (10). Moreover, wild-type FGFRs can promote tumorigenesis and resistance to both chemo- and targeted therapies, including KRAS inhibitors (11–15). Efforts to inhibit FGFRs have led to the development of FGFR-targeted therapies, which include selective, non-selective and covalent tyrosine kinase inhibitors, as well as monoclonal antibodies against the receptors and FGF ligand traps (10,16).

While the oncogenic functions of FGFRs in invasive cancers have been extensively studied, their role in the early phases of tumorigenesis remains largely unexplored. Here, we discovered that FGFR2 was up-regulated in mutant KRAS-driven pancreatic metaplasia and pre-neoplasia, supported PanIN formation and FGFR2 inactivation restricted PDA progression, making it a potential target for pancreatic cancer interception.

## RESULTS

### Analysis of matched organoid cultures reveals up-regulation of *Fgfr2* in pre-neoplastic compared to neoplastic KPC cells

To investigate the molecular mechanisms that promote PDA progression, we utilized the *Kras^LSLG12D/+^; Trp53^LSLR172H/+^; Pdx1-Cre* (KPC) mouse model in which *Kras*^G12D^ and *Trp53*^R172H^ were expressed in the pancreas by virtue of Cre recombinase under the control of the *Pdx1* promoter (17,18). In this genetically engineered mouse model (GEMM), the transition from pre-neoplasia to neoplasia is associated with loss of heterozygosity (LOH) of the wild-type *Trp53* allele (19). Indeed, we confirmed by PCR-based genotyping that loss of the wild-type *Trp53* allele occurred *in vivo* in neoplastic-enriched cell populations isolated from KPC tumors (Fig. 1A) (18,20). However, in all samples, we observed a faint band corresponding to the wild-type copy of *Trp53*, suggesting the presence of pre-neoplastic cells within the tumor.

**Figure 1.**
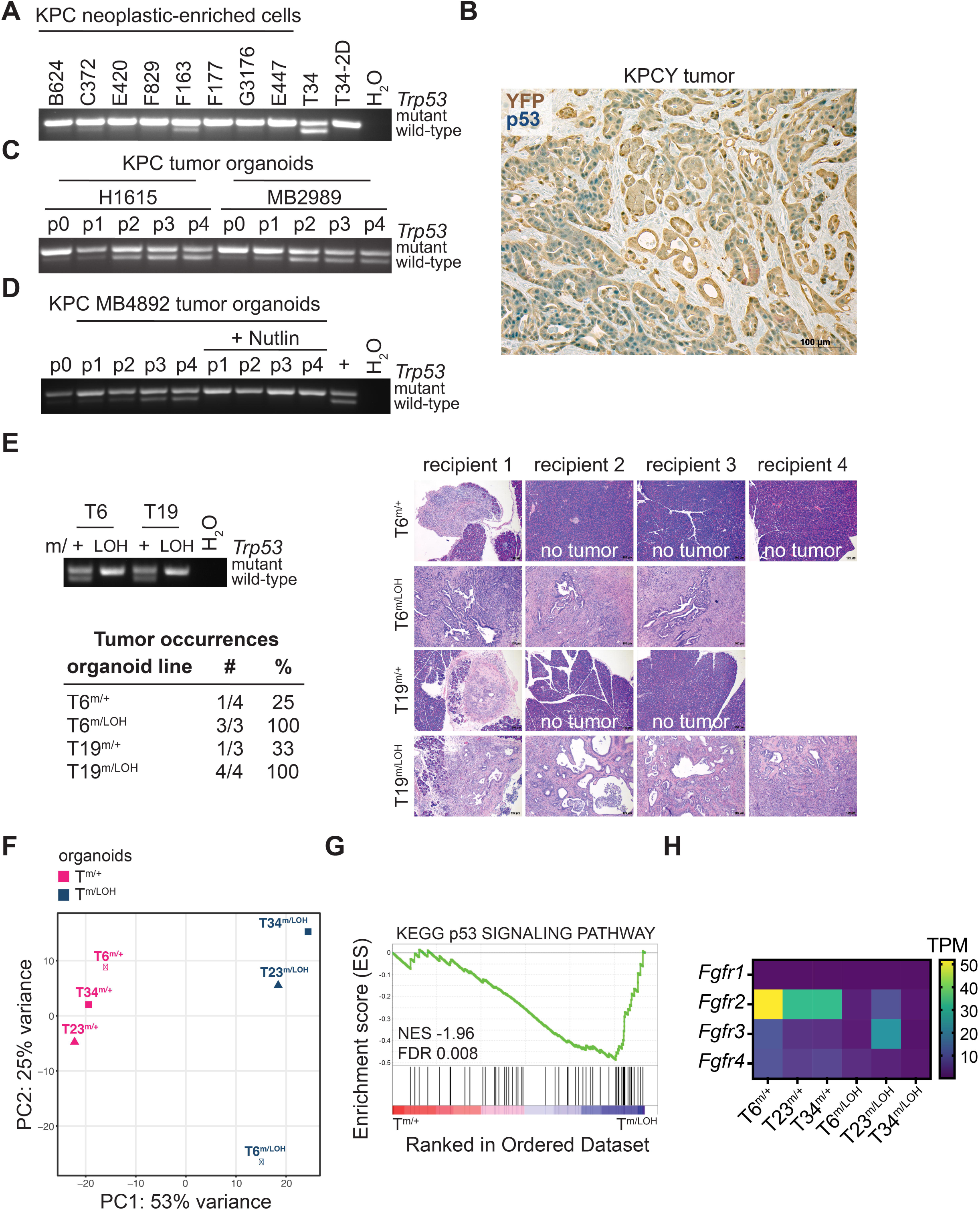
Analysis of matched organoid cultures reveals up-regulation of *Fgfr2* in pre-neoplastic compared to neoplastic KPC cells. **A.** PCR-based genotyping of neoplastic-enriched cells from KPC tumors for assessing LOH of the wild-type *Trp53* allele. Tumor ‘T34’ organoids and a 2D line derived from the same T organoids are analyzed as controls. **B.** Immunohistochemical staining for YFP (brown staining) and p53 (blue staining) in a KPCY tumor section. **C.** PCR-based genotyping for assessing the *Trp53* status in neoplastic-enriched cells from KPC tumors after isolation (p0) and upon passaging as organoids (p1 to p4). **D.** PCR-based genotyping for assessing the *Trp53* status in neoplastic-enriched cells from a KPC tumor after isolation (p0) and upon passaging as organoids (p1 to p4) in medium with or without Nutlin. +, positive PCR control **E.** T^m/LOH^ organoids are more aggressive *in vivo* than T^m/+^ organoids. The *Trp53* status of the organoid lines transplanted was assessed by PCR-based genotyping. Table of primary tumor occurrences. Hematoxylin and eosin (H&E) staining of tumors or pancreata (if no tumor was observed). Scale bars: 100 μm **F.** PCA of RNA-seq data of T organoid pairs (n = 3). Samples are colored based on their group: T^m/+^ in pink and T^m/LOH^ organoids in blue. **G.** GSEA signature ‘KEGG p53 SIGNALING PATHWAY’ is repressed in T^m/LOH^ compared to T^m/+^ organoids. NES, normalized enrichment score; FDR, false discovery rate. **H.** *Fgfrs* expression as TPM in T^m/+^ and T^m/LOH^ organoids as determined by RNA-seq.

To investigate if the tumor mass contained pre-neoplastic cells, we analyzed p53 expression in tumor sections of KPCY mice (*Kras^LSLG12D/+^; Trp53^LSLR172H/+^; Pdx1-Cre; Rosa26^LSLYFP^*), in which all *Kras*^G12D^-expressing cells presented the yellow fluorescent protein (YFP) lineage label (21). Under basal conditions, p53 levels are undetectable by immunohistochemistry (IHC) due to its rapid turnover (18,22,23). Mutations in *Trp53* result in highly stabilized mutant p53 proteins only in the absence of the other wild-type *Trp53* allele (18,19,24). Therefore, only *Trp53* LOH cells present positive staining for p53 by IHC. Immunohistochemical labeling for YFP and p53 in KPCY tumors revealed two populations of YFP-positive cells: one with increased nuclear expression of p53 which represented *Trp53* LOH neoplastic cells, and the other in which p53 was not detected, representing the pre-neoplastic cells (Fig. 1B). Indeed, cells that did not present p53 accumulation displayed histological features of acinar-to-ductal metaplasia (ADM) and murine PanINs (mPanINs); while cells with detectable p53 expression exhibited features of invasive carcinoma.

We previously described the establishment of pancreatic ductal organoid cultures from multiple primary tumors from KPC mice and showed that tumor ‘T’ organoids did not exhibit *Trp53* LOH (25). Given that *Trp53* LOH was prevalent in freshly isolated KPC neoplastic cells (Fig. 1A), we sought to determine when *Trp53* LOH cells were lost in organoid culture. We performed PCR-based genotyping of neoplastic-enriched cell populations after isolation and upon passaging as organoids (Fig. 1C). Pre-neoplastic cells harboring both mutant and wild-type *Trp53* alleles outcompeted the *Trp53* LOH cells in organoid culture in a few passages. This growth advantage was independent of medium or matrix composition and associated with the ploidy status (Supplementary Fig. S1A-C). KPC tumors were enriched in aneuploid and tetraploid cells which were outcompeted in organoid culture by diploid and quasi-diploid cells.

To isolate the *Trp53* LOH cells, we treated the organoids with Nutlin, a small molecule that mediates p53 stabilization by inhibiting the p53-MDM2 interaction (26). Nutlin treatment induced the death of the cells that retained wild-type p53 and allowed for the isolation of the *Trp53* LOH cells (Fig. 1D; Supplementary Fig. S1C-E). Using this strategy, we established matched organoid cultures of KPC pre-neoplastic cells that retained the wild-type *Trp53* allele (T^m/+^) and KPC neoplastic cells that had lost the wild-type *Trp53* allele (T^m/LOH^).

Once stable cultures were established, T^m/+^ organoids were diploid, while some T^m/LOH^ organoids showed a slight increase in DNA content by Propidium Iodide (PI) staining (Supplementary Fig. S2A) (25). While T^m/+^ and T^m/LOH^ organoids proliferated at similar rate *in vitro*, T^m/LOH^ organoids formed tumors with significantly higher penetrance following orthotopic implantation in syngeneic mice (Fig. 1E; Supplementary Fig. S2B). Mice transplanted with T^m/+^ organoids developed either no tumor or very small tumors when pancreata were collected at the humane endpoint of mice transplanted with the matched T^m/LOH^ organoids.

To investigate the transcriptional differences between KPC pre-neoplastic and neoplastic cells, we performed RNA sequencing (RNA-seq). When subjected to principal component analysis (PCA), T^m/+^ and T^m/LOH^ organoids separated based on their LOH status and paired organoids did not cluster together (Fig. 1F; Supplementary Fig. S2C). Comparison of T^m/+^ and T^m/LOH^ organoids identified 1514 differentially expressed genes: 996 up-regulated and 518 down-regulated genes in T^m/LOH^ relative to T^m/+^ (Supplementary Table S1). Gene Set Enrichment Analysis (GSEA) confirmed repression of the p53 signaling pathway in T^m/LOH^ organoids (Fig. 1G). Among the up-regulated genes, we identified several genes that were previously associated with pancreatic cancer progression, such as *Prdm1/*BLIMP1, *Foxa1, Gata5*, *Twist1, Glul* and *Soat1* (Supplementary Fig. S2D) (20,27–30). Among the genes induced in T^m/+^ compared to T^m/LOH^ organoids, we noted *Fgfr2* isoform IIIb (from now on, *Fgfr2*), which we had previously shown to be expressed in KPC pre-neoplastic cells (Fig. 1H; Supplementary Fig. S2E) (31). Of note, *Fgfr2* was the only *Fgfr* to be highly expressed in T^m/+^ organoids.

### FGFR2 is progressively up-regulated in mutant KRAS-driven pancreatic metaplasia, pre-neoplasia and human Classical PDA

To investigate FGFR2 expression during pancreatic cancer progression, we performed immunofluorescent labeling (IF). In healthy pancreata, FGFR2 was not expressed in normal ducts or acinar cells, but was weakly expressed in blood vessels (Fig. 2A). Notably, the percentage of acinar cells that expressed FGFR2 significantly increased as soon as KRAS^G12D^ initiated the transformation of these cells to metaplastic ductal-like lesions in young *Kras^LSLG12D/+^; Pdx1-Cre* (KC) mice. In these mice, the majority of cytokeratin 19 (CK19)-positive pre-neoplastic cells expressed FGFR2 at the plasma membrane (Fig. 2B). In KPC tumors, FGFR2 expression was observed in pre-neoplastic cells in which p53 was not or barely detectable, whereas it was absent or reduced in invasive cells with nuclear accumulation of p53 (Fig. 2C) (31).

**Figure 2.**
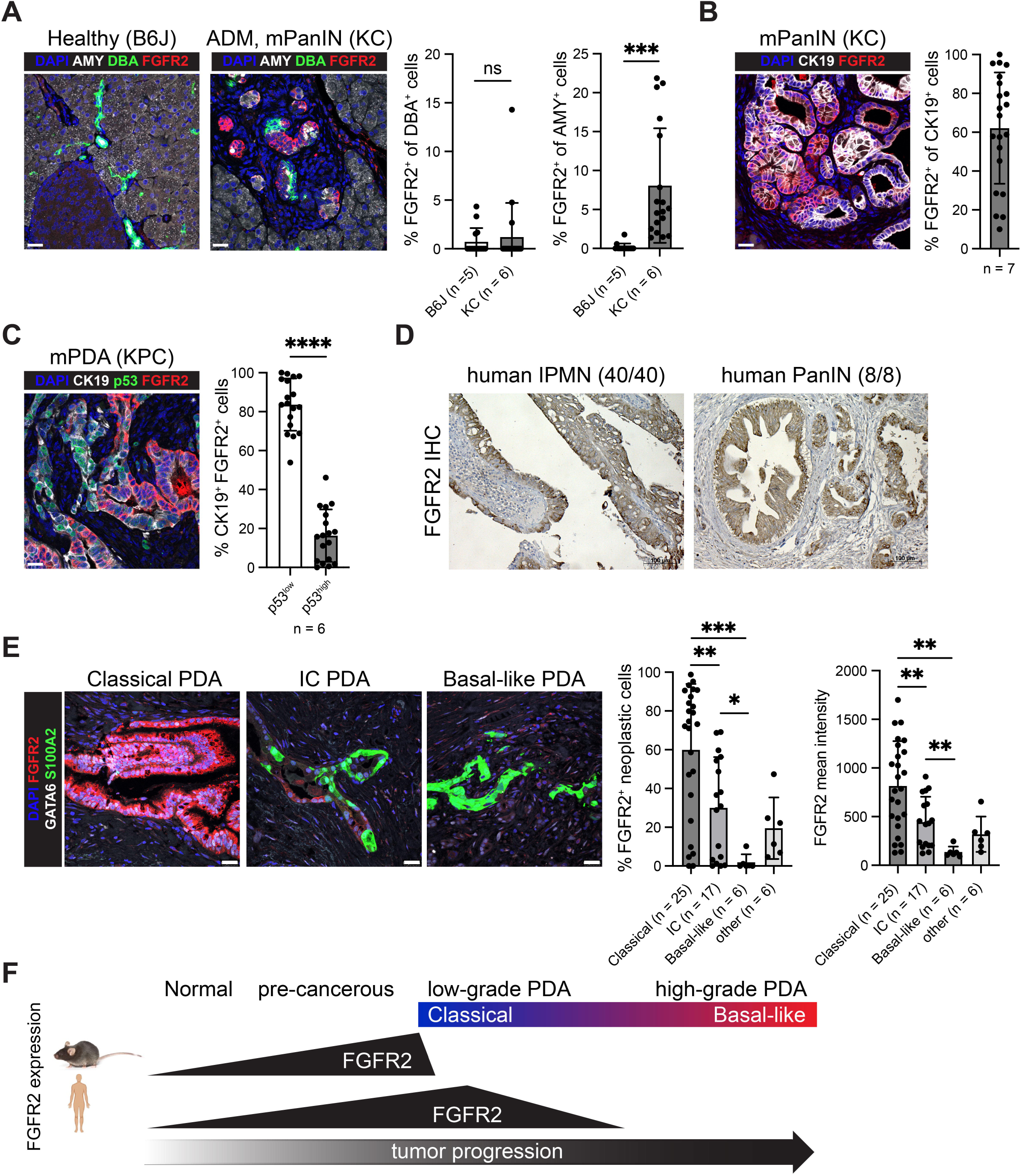
FGFR2 is progressively up-regulated in mutant KRAS-driven pancreatic metaplasia, pre-neoplasia and human Classical PDA. **A.** Left, representative IF for the acinar marker AMY (white), the ductal marker Dolichos Biflorus Agglutinin (DBA) (green), FGFR2 (red) and 4′,6-diamidino-2-phenylindole (DAPI, blue) conducted on healthy pancreata from B6J mice (n = 5) and ADM and mPanINs from KC mice (n = 6). Scale bars, 25 μm. Right, quantification of staining plotted as mean ± SD. 3 images per mouse were quantified. Unpaired Student’s t test. **B.** Left, representative IF for CK19 (white), FGFR2 (red) and DAPI (blue) conducted on mPanINs from KC mice (n = 7). Scale bars, 25 μm. Right, quantification of staining plotted as mean ± SD. 3 images per mouse were quantified. **C.** Left, representative IF for CK19 (white), p53 (green), FGFR2 (red) and DAPI (blue) conducted on tumors from KPC mice (n = 6). Scale bars, 25 μm. Right, quantification of staining plotted as mean ± SD. 3 images per mouse were quantified. Unpaired Student’s t test. **D.** Representative IHC for FGFR2 conducted on human IPMN and PanIN lesions. **E.** Left, representative IF for FGFR2 (red), GATA6 (white), S100A2 (green) and DAPI (blue) conducted on human PDAs. Scale bars, 25 μm. Samples that present % GATA6^+^ S100A2^-^ cancerous cells > 60 are classified as ‘Classical’, samples with % GATA6^-^ S100A2^+^ cancerous cells > 30 are classified as ‘Basal-like’ and samples with % GATA6^+^ S100A2^+^ cancerous cells > 5 are classified as ‘IC’. Remaining samples that don’t meet these criteria are classified as ‘other’. Right, quantification of staining plotted as mean ± SD. Unpaired Student’s t test. **F.** Schematic representation of FGFR2 expression during pancreatic cancer progression in the KRAS^G12D^-driven mouse model and in human samples.

FGFs are produced by fibroblasts and bind to heparin and heparan sulfates, which protect them from proteases and enhance the formation of an active signaling complex with the FGFRs (32). We found that the FGFR2’s ligands FGF7 and FGF10 often colocalized with FGFR2 and the percentage of double positive cells increased with disease progression, potentially due to the rise in fibroblast numbers (Supplementary Fig. S3A, B) (15).

Next, we examined whether the expression pattern of FGFR2 observed in mice was conserved in human. Consistent with the finding in mice, human precursor lesions intraductal papillary mucinous neoplasm (IPMN) and PanIN expressed FGFR2 (Fig. 2D). However, differently from mice, FGFR2 was expressed at high levels in most neoplastic cells in Classical PDAs, in fewer cells and at lower levels in Intermediate Co-expressor (IC) PDAs, and was absent in Basal-like PDAs, as defined by immunolabeling for the Classical marker GATA6 and the Basal-like marker S100A2 (Fig. 2E) (33). In line with these results, analysis of expression data of human PDAs showed significantly higher expression of *FGFR2* in Classical compared to Basal-like PDAs (Supplementary Fig. S3C).

Since GATA6 and FGFR2 were highly expressed in Classical PDAs, we wondered whether GATA6 regulated *FGFR2* expression. In T^m/+^ organoids, *Gata6* knock-down using 3 different doxycycline-inducible short hairpin RNAs resulted in *Fgfr2* down-regulation (Supplementary Fig. S3D). In addition, CUT&RUN profiling in the Classical PDA cell line HPAF-II revealed the binding of GATA6 to the promoter region of *FGFR2* (31). Together these results demonstrated that FGFR2 is progressively up-regulated in human pancreatic pre-neoplasia and Classical PDA and repressed in Basal-like PDA (Fig. 2F), and that its expression is regulated by GATA6.

### FGFR2 is repressed by TGFβ signaling

Increased activation of the TGFβ pathway has been reported in invasive KPC cells and in Basal-like PDAs, where it is known to contribute to epithelial-to-mesenchymal transition (EMT) (31,34,35). Since FGFRs are also key regulators of EMT during embryonic development and in neoplastic cells during cancer progression (36), we sought to evaluate whether TGFβ signaling mediates FGFR2 repression, as previously observed in mammary epithelial cells (37). Therefore, we treated 3 T^m/+^ organoid lines with TGFβ and analyzed the transcriptional changes by RT–qPCR (Fig. 3A). Organoids cultured in ‘Complete’ medium expressed *Fgfr2* at higher levels compared to organoids grown in ‘Minimal’ medium without any additives including the TGFβ receptor inhibitor A83-01 and the FGFR2 ligand FGF10. The addition of TGFβ to the ‘Minimal’ medium further reduced the expression of *Fgfr2* and induced the up-regulation of the mesenchymal genes *Fgfr1, Snai1* and *Fn1*. The switch from *Fgfr2* to *Fgfr1* expression was dose-dependent and time-dependent (Fig. 3B; Supplementary Fig. S4A).

**Figure 3.**
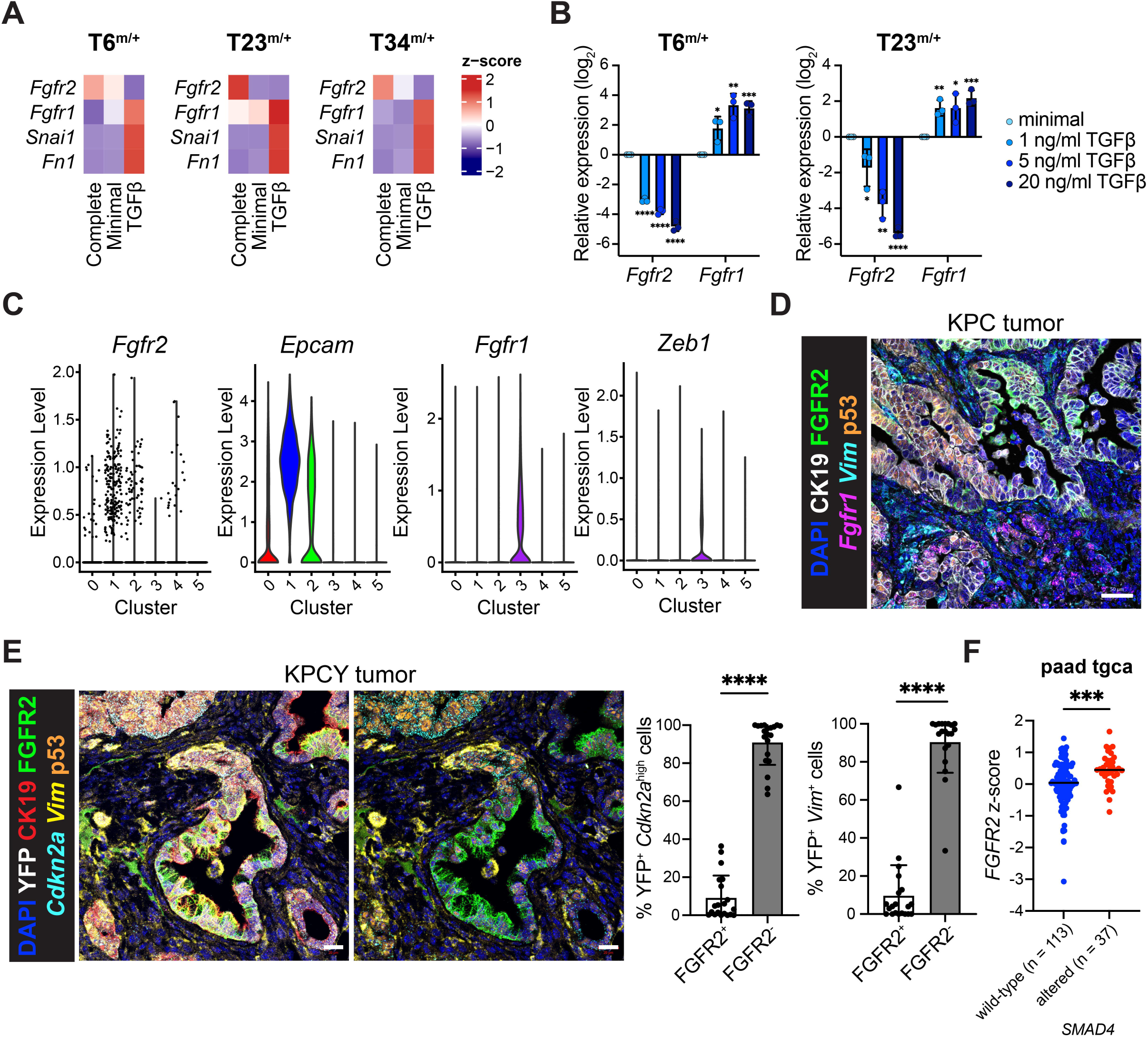
FGFR2 is repressed by TGFβ signaling. **A.** Heatmap of average z-score values of *Fgfr2, Fgfr1, Snai1* and *Fn1* expression as determined by RT-qPCR in T^m/+^ organoids after growth in Complete organoid medium, in Minimal medium or Minimal medium supplemented with TGFβ (n = 3). **B.** *Fgfr2* and *Fgfr1* expression as determined by RT-qPCR in T6^m/+^ and T23^m/+^ organoids after growth in Minimal medium supplemented with different doses of TGFβ or vehicle (n = 3). Results show mean ± SD. Unpaired Student’s t test. **C.** Violin plots showing the expression of *Fgfr2, Epcam, Fgfr1* and *Zeb1* in the different populations of KPC pre-neoplastic and neoplastic cells (31). **D.** Representative RNA ISH of *Fgfr1* (magenta), *Vim* (cyan) combined with IF for CK19 (white), FGFR2 (green), p53 (orange) and DAPI (blue) in tumors from KPC mice. Scale bar, 50 μm. **E.** Left, representative RNA ISH of *Cdkn2a/*ARF (cyan), *Vim* (yellow) combined with IF for YFP (white), CK19 (red), FGFR2 (green), p53 (orange) and DAPI (blue) conducted on tumors from KPCY mice (n = 5). Scale bar, 20 μm. Right, quantification of staining plotted as mean ± SD. 4 images per mouse were quantified. Unpaired Student’s t test. **F.** *FGFR2* z-score values in *SMAD4* wild-type and altered PDA samples (77). Results show mean ± SD. Unpaired Student’s t test.

To assess *Fgfr2* and *Fgfr1* expression in relation to cell states, we analyzed our scRNA-seq data of KPC pre-neoplastic and neoplastic cells (Fig. 3C) (31). *Fgfr2* expression was detected in few of the epithelial cells in cluster 1, while *Fgfr1* was expressed by the mesenchymal cells in cluster 3. Using RNA *in situ* hybridization (RNA ISH) in combination with IF for p53 and *Cdkn2a/*ARF as markers of invasive cells, *Vimentin* as a marker of mesenchymal cells and CK19 as a marker of ductal cells, we confirmed that FGFR2 was expressed in pre-neoplastic ductal lesions in murine tumors, while *Fgfr1* was expressed in invasive mesenchymal cells and some stromal cells (Fig. 3D, E; Supplementary Fig. S4B, C). Although PDA cells occupy a continuum of epithelial-to-mesenchymal expression states (31,34), *Fgfr2* expression was restricted to pre-neoplastic cells with strong epithelial features, while *Fgfr1* was uniquely expressed by mesenchymal cells. The majority of neoplastic cells that exhibited a partial EMT phenotype expressed neither gene.

Finally, we found that *FGFR2* expression was significantly increased in *SMAD4* altered human PDAs compared with those expressing wild-type *SMAD4*, further supporting a role for TGFβ signaling in down-regulating *FGFR2* (Fig. 3F).

### FGFR2 is dispensable for pancreas recovery following injury

FGFR2 up-regulation in mutant KRAS-driven pancreatic metaplasia and pre-neoplasia could be part of a tumor-promoting program or an inflammatory response. Since FGFR2 was implicated in the repair of the skin, intestine, liver and lung (38), we tested if FGFR2 was required for pancreas recovery following injury. We inactivated *Fgfr2* in the pancreas by crossing *Pdx1-Cre* (C) to *Fgfr2^f/f^* mice in which the alternatively spliced FGF binding Ig domain IIIb, IIIc and transmembrane domain of *Fgfr2* are flanked by loxP sites, resulting in an inactive protein after recombination (39) (Supplementary Fig. S5A, B). The pancreata of C *Fgfr2^f/f^* mice did not present any overt defects upon histological evaluation (Fig. 4A). Using an established model of pancreatitis with the cholecystokinin analog cerulein, we found that one day after treatment the pancreata of both C *Fgfr2^+/+^* and *Fgfr2^f/f^*mice presented transient pancreatic inflammation, with edema, acinar cell damage, ADM and infiltration of immune cells. By day 7, the tissue integrity in both strains was almost completely restored, reaching full recovery by day 30. Immunolabeling was consistent with the histological analysis, revealing a transient decrease of the acinar marker amylase and concomitant increase of the ductal marker CK19 at day 1 and reversion to pre–cerulein treatment levels by day 30 in both C *Fgfr2^+/+^* and *Fgfr2^f/f^* mice (Fig. 4B). These results suggested that *Fgfr2* is dispensable for pancreas recovery following injury.

**Figure 4.**
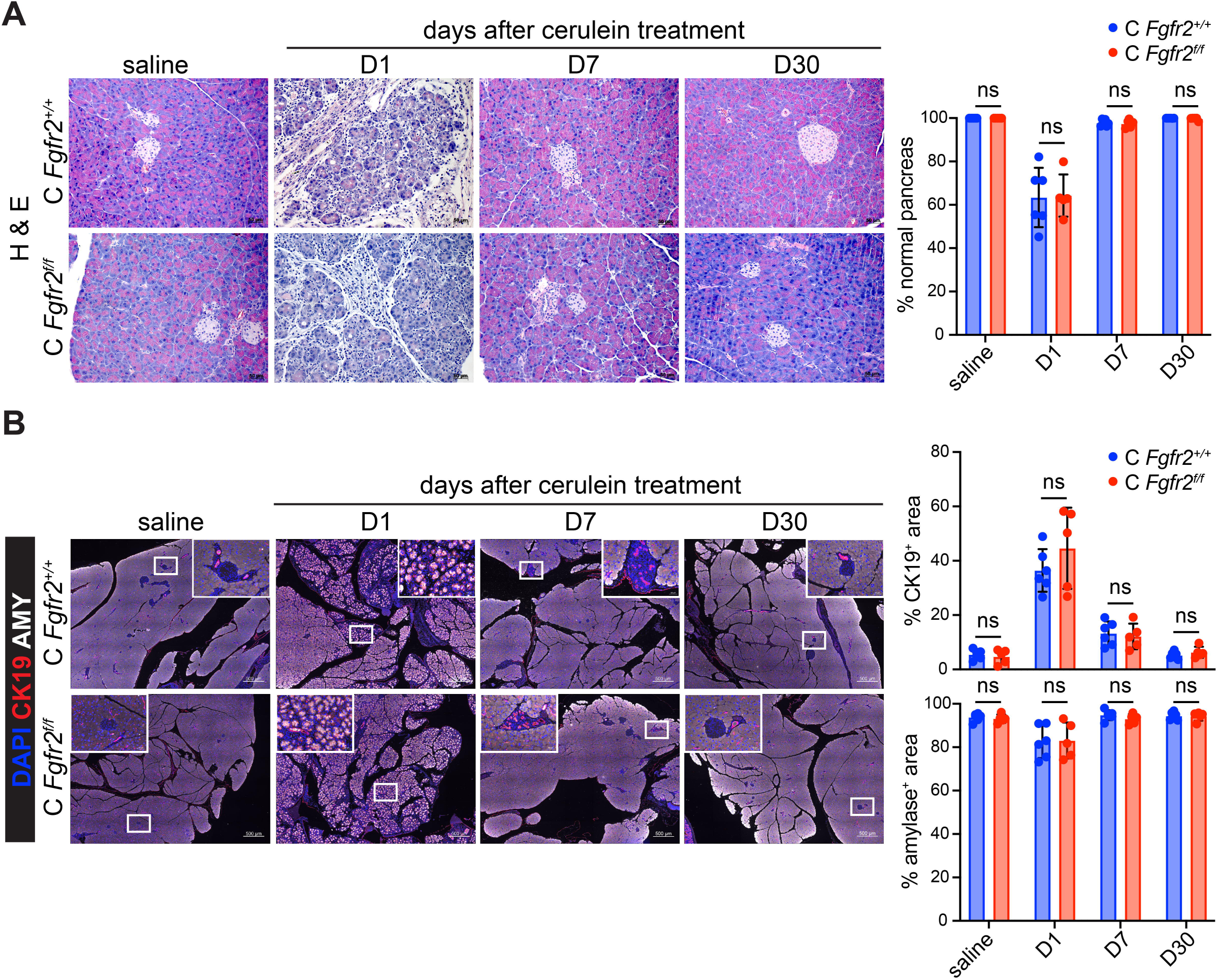
FGFR2 is dispensable for pancreas recovery following injury. **A.** Left, representative H&E staining conducted on pancreatic samples from C *Fgfr2^+/+^* and *Fgfr2^f/f^* mice at different time points after saline or cerulein treatment (day −1, day 0). Scale bar, 50 μm. Right, quantification of the fraction of normal pancreas (n = 5-6). Results show mean ± SD. Unpaired Student’s t test. **B.** Left, representative IF for CK19 (red), amylase (white) and DAPI (blue) conducted on pancreatic samples from C *Fgfr2^+/+^*and *Fgfr2^f/f^* mice at different time points after saline or cerulein treatment (day −1, day 0) (n = 5-6). Scale bars are 500 μm for main images and 50 μm for insets. Right, quantification of staining plotted as mean ± SD. Unpaired Student’s t test.

### FGFR2 facilitates mutant KRAS-induced transformation of acinar cells by promoting proliferation and MAPK pathway activation

Cerulein-mediated injury to the pancreas accelerates mutant KRAS-driven metaplastic lesions and mPanIN formation (40), as confirmed by histological and immunolabeling analyses of the pancreata from KC in comparison to C mice (Supplementary Fig. S6A, B). FGFR2 expression was not detected by IHC upon transient pancreatitis and following recovery (Fig. 5A). However, in KC mice, FGFR2 was up-regulated in acinar cells, metaplastic lesions and mPanINs upon KRAS^G12D^-driven malignant transformation, indicating its potential involvement in promoting early progression. Intriguingly, analysis of normal adjacent tissue of PDA patients revealed areas of morphologically normal acinar cells, ADM and PanINs that stained positive for FGFR2, which was reminiscent of what we observed in KC mice (Supplementary Fig. S6C).

**Figure 5.**
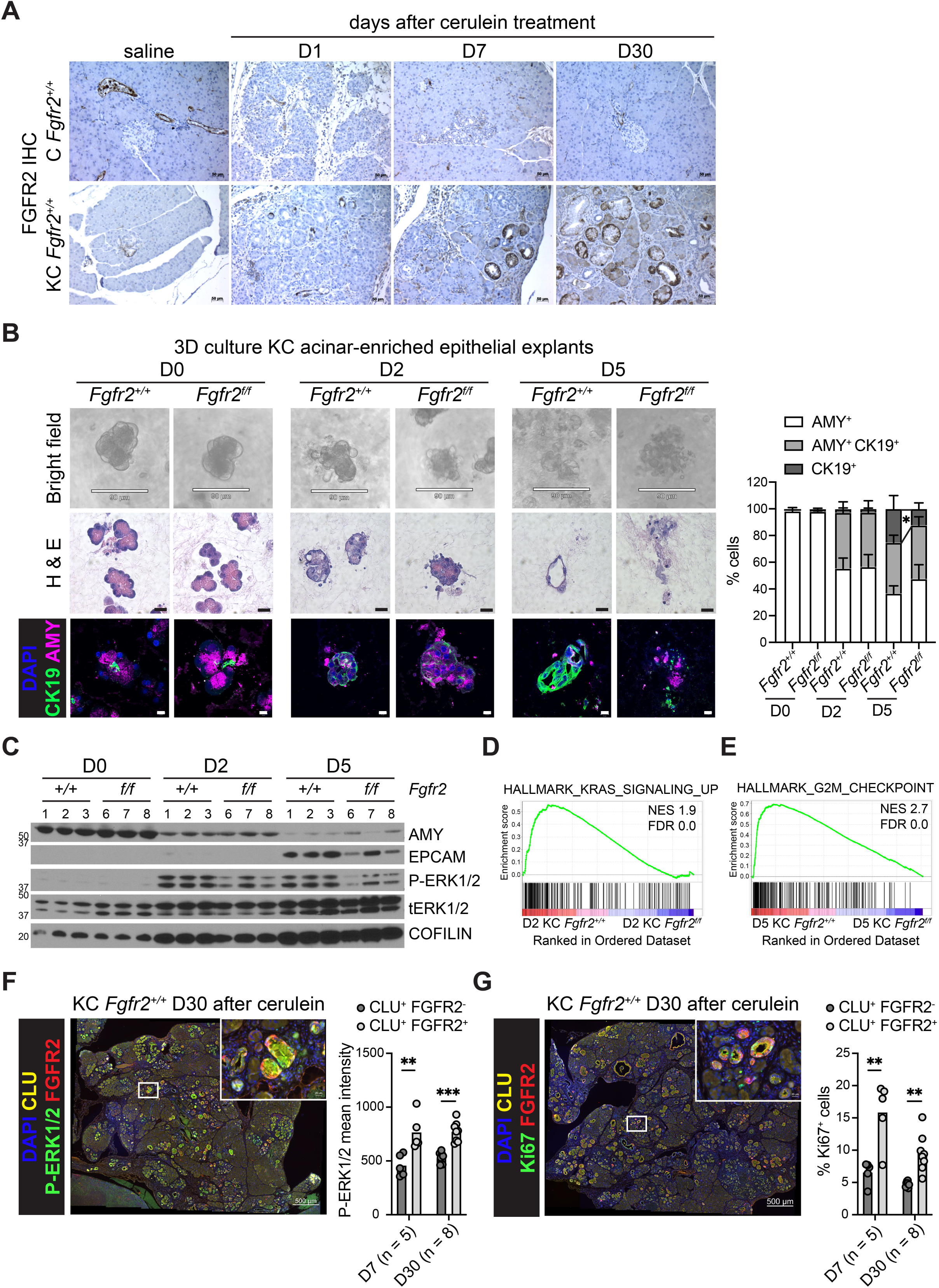
FGFR2 facilitates mutant KRAS-induced transformation of acinar cells by promoting proliferation and MAPK pathway activation. **A.** Representative IHC for FGFR2 conducted on pancreatic samples from C and KC mice at different time points after saline or cerulein treatment (day −1, day 0). Scale bar, 50 μm. **B.** Left, representative bright field images (scale bar, 90 μm), H&E staining (scale bar, 20 μm) and IF for CK19 (green), amylase (magenta) and DAPI (blue) (scale bar, 10 μm) conducted on acinar clusters and associated terminal ductal epithelium from KC *Fgfr2^+/+^* and *Fgfr2^f/f^*mice at different time points after culture in collagen. Right, quantification of staining plotted as mean ± SD (n = 5). Unpaired Student’s t test. **C.** Protein expression analysis in acinar clusters and associated terminal ductal epithelium from KC *Fgfr2^+/+^* and *Fgfr2^f/f^*mice at different time points after culture in collagen, as determined by Western blotting. Loading control, COFILIN. **D.** GSEA signature ‘HALLMARK KRAS SIGNALING UP’ is increased in acinar-enriched explants from KC *Fgfr2^+/+^*compared to *Fgfr2^f/f^* mice at day 2 of culture in collagen. NES, normalized enrichment score; FDR, false discovery rate. **E.** GSEA signature ‘HALLMARK G2M CHECKPOINT’ is increased in acinar-enriched explants from KC *Fgfr2^+/+^* compared to *Fgfr2^f/f^* mice at day 5 of culture in collagen. NES, normalized enrichment score; FDR, false discovery rate. **F, G.** Left, representative IF for CLU (yellow), FGFR2 (red), phospho-ERK1/2 (**F**) or Ki67 (**G**) (green), and DAPI (blue) conducted on pancreatic samples from KC *Fgfr2^+/+^* mice at day 7 and 30 after cerulein treatment (day −1, day 0) (n = 5-8). Scale bars are 500 μm for main images and 20 μm for insets. Right, quantification of staining plotted as mean ± SD. Paired Student’s t test.

To explore the role of FGFR2 in pancreatic tumorigenesis, we crossed KC to *Fgfr2^f/f^* mice (Supplementary Fig. S6D). Next, we modeled mutant KRAS-induced malignant transformation of acinar cells using 3D cultures of KC acinar-enriched epithelial explants embedded in collagen (41). Under these culture conditions, many acinar cells were lost because of cell death, but some acinar clusters and associated ductal cells formed ductal structures (Fig. 5B). We found that acinar-enriched clusters from KC *Fgfr2^+/+^* mice formed ductal structures more efficiently than the ones from KC *Fgfr2^f/f^*mice. Protein analysis at day 0, 2 and 5 of culture showed a progressive decreased expression of the acinar marker amylase, and conversely an increased expression of the ductal marker EPCAM; however, EPCAM up-regulation was greater in KC *Fgfr2^+/+^* compared to KC *Fgfr2^f/f^* cells (Fig. 5C; Supplementary Fig. S6E). Furthermore, analysis of MAPK signaling by detection of the phosphorylation of ERK1/2 revealed activation of the pathway from day 2 of culture, which was stronger in KC *Fgfr2^+/+^* compared to KC *Fgfr2^f/f^*cells (Fig. 5C; Supplementary Fig. S6E). mRNA expression analysis by RNA-seq showed progressive repression of acinar genes and up-regulation of ductal genes, including *Fgfr2*, over time in culture (Supplementary Fig. S6F, Table S2). GSEA revealed up-regulation of KRAS signaling at day 2 and proliferation-associated programs, such as G2M checkpoint, E2F and MYC targets, at day 5 in KC *Fgfr2^+/+^* compared to KC *Fgfr2^f/f^* cells (Fig. 5D, E; Supplementary Fig. S6G, Table S2). Altogether, these results suggested that FGFR2 promotes proliferation and enhances MAPK signaling during mutant KRAS-driven acinar-to-ductal-like cell transformation *ex vivo*. To corroborate these findings, we performed IF for FGFR2, Ki67, phospho-ERK1/2 and Clusterin (CLU), a marker of acinar transformation (42), in pancreata from KC mice at day 7 and 30 after cerulein treatment. We found that CLU- and FGFR2-positive cells presented with a stronger signal for phospho-ERK1/2, and a higher fraction of these cells were Ki67-positive (Fig. 5F, G), further reinforcing that FGFR2 supports mutant KRAS-induced malignant transformation of acinar cells by promoting proliferation and MAPK signaling activation. This suggested that FGFR2 is a key factor in pushing KRAS signaling over the threshold necessary for disease progression.

### FGFR2 promotes KRAS^G12D^-driven pancreatic tumorigenesis

To assess the effect of FGFR2 inactivation on tumorigenesis *in vivo*, we analyzed metaplastic lesion and mPanIN formation during spontaneous and cerulein-accelerated tumorigenesis in KC *Fgfr2^f/f^*compared to *Fgfr2^+/+^* mice. In both models, we found that KC *Fgfr2^+/+^* mice presented a significantly larger fraction of diseased pancreatic tissue compared to age-matched KC *Fgfr2^f/f^* mice (Fig. 6A, B, D, E), indicating that mutant KRAS-driven pancreatic tumorigenesis was hindered upon inactivation of FGFR2. Immunolabeling was consistent with the histological analysis, revealing smaller areas of mPanINs and larger areas of acinar tissue in KC *Fgfr2^f/f^* compared to *Fgfr2^+/+^* mice (Supplementary Fig. S7A, B).

**Figure 6.**
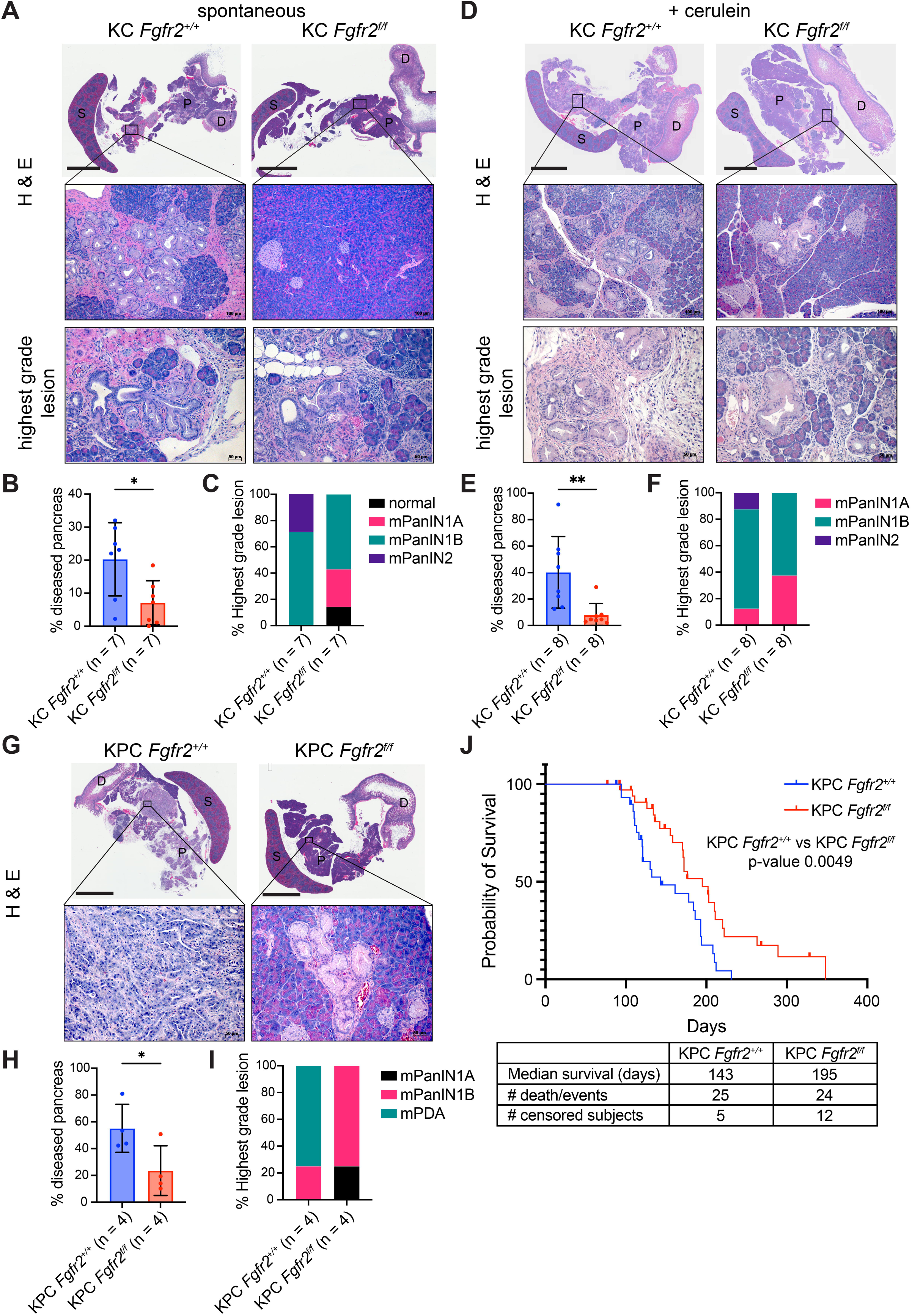
FGFR2 promotes KRAS^G12D^-driven pancreatic tumorigenesis. **A, D.** Representative H&E staining of the pancreata and highest-grade lesions from 7- to 11- months-old KC *Fgfr2^+/+^* and *Fgfr2^f/f^* mice (n = 7) (**A**) or from KC *Fgfr2^+/+^* and *Fgfr2^f/f^* mice 30 days after cerulein treatment (day −1, day 0) (n = 8) (**D**). Scale bars are 4 mm for pancreas images, 100 μm for insets and 50 μm for highest grade lesions images. S, spleen; D, duodenum; P, pancreas **B, E.** Quantification of the fraction of diseased pancreas. Results show mean ± SD. Unpaired Student’s t test. **C, F.** Classification of highest-grade lesions. **G.** Representative H&E staining of the pancreata and highest-grade lesions from 4-months-old KPC *Fgfr2^+/+^*and *Fgfr2^f/f^* mice (n = 4). Scale bars are 5 mm for pancreas images and 50 μm for insets. S, spleen; D, duodenum; P, pancreas **H.** Quantification of the fraction of diseased pancreas. Results show mean ± SD. Unpaired Student’s t test. **I.** Classification of highest-grade lesions. **J.** Kaplan–Meier survival curve of percent survival for KPC *Fgfr2^+/+^* (n = 25) and KPC Fgfr2^f/f^ (n = 24) mice. Notches represent enrolled mice that died from causes other than tumor (i.e., lymphoma, papilloma). Log-rank Mantel–Cox test. Table of median survival (days).

In addition to having lower fraction of diseased pancreas, KC *Fgfr2^f/f^*mice harbored lower grade mPanINs. In the spontaneous tumorigenesis model, the KC *Fgfr2^+/+^* mice exhibited mPanIN-1B and mPanIN-2 lesions, while KC *Fgfr2^f/f^* mice harbored mPanIN-1A, mPanIN-1B or no lesions (Fig. 6C). Similarly, in the cerulein-accelerated tumorigenesis model, the KC *Fgfr2^+/+^*mice exhibited mPanIN-1A, mPanIN-1B and mPanIN-2 lesions, while KC *Fgfr2^f/f^*mice harbored the lower grade mPanIN-1A and mPanIN-1B (Fig. 6F).

Given the differences that we observed in pre-neoplastic lesion formation in the KC pancreatic cancer model with *Fgfr2* inactivation, we evaluated tumor formation in the KPC model. At 4 months of age, KPC *Fgfr2^+/+^*mice presented a significantly larger fraction of diseased pancreas tissue compared to KPC *Fgfr2^f/f^* mice (Fig. 6G, H). Notably, 3 out of 4 of the KPC *Fgfr2^+/+^* mice examined had developed a tumor mass, while KPC *Fgfr2^f/f^* mice harbored mPanIN-1A or mPanIN-1B lesions (Fig. 6I). Next, we followed a cohort of mice to humane endpoint and found that *Fgfr2* inactivation in the KPC model led to a significant extension of median survival from 143 to 195 days in KPC *Fgfr2^f/f^,* although KPC *Fgfr2^f/f^*mice eventually all succumbed to malignant disease (Fig. 6J). Genotyping analysis of organoids derived from KPC *Fgfr2^f/f^* tumors showed recombination of the *Kras, Trp53* and *Fgfr2* locus, thus excluding the possibility of incomplete recombination of *Fgfr2* as a potential escape mechanism (Supplementary Fig. S7C). Immunolabeling analysis revealed similar levels of phosphorylated ERK1/2 and percentage of proliferating pre-neoplastic cells in mPanINs from KC *Fgfr2^+/+^* and *Fgfr2^f/f^*mice (Supplementary Fig. S7D-G). Altogether, these data demonstrated that abrogation of FGFR2 delays KRAS^G12D^- induced tumorigenesis, with alternative adaptative pathways likely compensating for the inactivation of FGFR2 later in tumor progression.

### FGFR2 and EGFR cooperate to promote mutant KRAS-driven pancreatic tumorigenesis

Previously, other groups showed that EGFR signaling is required for mutant KRAS-driven pancreatic cancer development (4,5). However, given that pharmacological inhibition of EGFR restricted but did not abrogate pre-neoplastic lesion formation (4,43), we wondered whether EGFR and FGFR2 cooperated in promoting pancreatic tumorigenesis. We first examined FGFR2 expression and function upon EGFR inhibition. Treatment of T^m/+^ organoids with the EGFR inhibitors gefitinib and erlotinib resulted in FGFR2 protein up-regulation, suggesting that FGFR2 may compensate for EGFR inhibition (Fig. 7A). Indeed, the growth of *Fgfr2* knock-out T^m/+^ organoids was strongly inhibited by erlotinib, while it was not consistently reduced compared to control organoids in normal culture conditions (Fig. 7B; Supplementary Fig. S8A). Overexpression of FGFR2 in T^m/+^ *Fgfr2^f/f^* organoids rescued the growth inhibition caused by erlotinib treatment (Fig. 7C).

**Figure 7.**
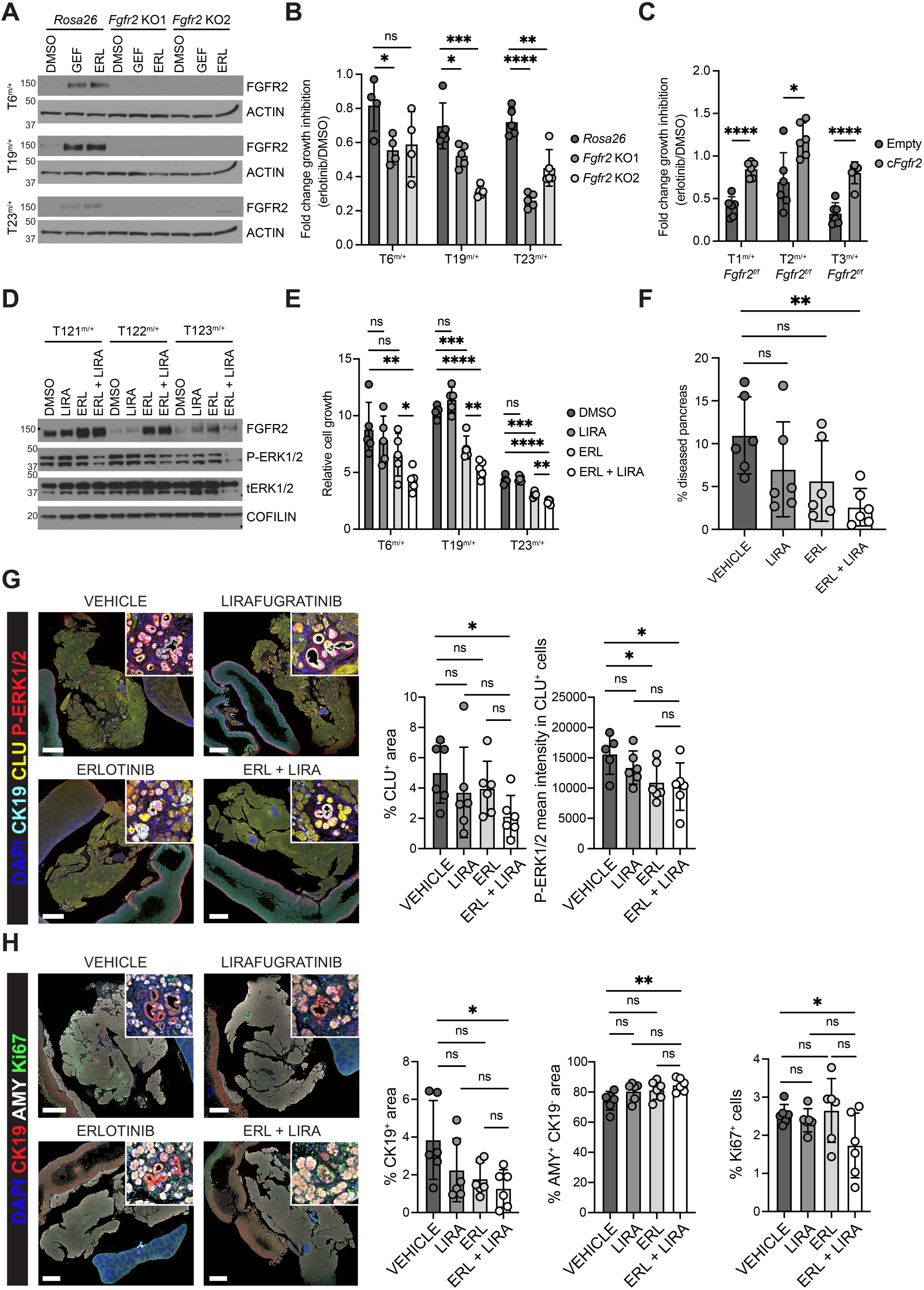
FGFR2 and EGFR cooperate to promote mutant KRAS-driven pancreatic tumorigenesis. **A.** FGFR2 protein expression in T^m/+^ organoids knocked-out for *Fgfr2* or control *Rosa26* treated with vehicle, gefitinib or erlotinib as determined by Western blotting. Loading control, ACTIN. **B.** Fold change of growth inhibition upon erlotinib treatment of T^m/+^ organoids knocked-out for *Fgfr2* or control *Rosa26* (n = 5). Results show mean ± SD. Unpaired Student’s t test. **C.** Fold change of growth inhibition upon erlotinib treatment of T^m/+^ *Fgfr2^f/f^* organoids expressing *Fgfr2* cDNA or control (n = 6). Results show mean ± SD. Unpaired Student’s t test. **D.** Protein expression analysis of T^m/+^ organoids treated with vehicle, lirafugratinib, erlotinib or the combination as determined by Western blotting. Loading control, COFILIN. **E.** Relative growth (day 4 / day 1) of T^m/+^ organoids treated with vehicle, erlotinib, lirafugratinib or the combination (n = 5). Results show mean ± SD. Unpaired Student’s t test. **F.** Quantification of the fraction of diseased pancreas in 2-months-old KC mice treated with cerulein for 2 days and the indicated inhibitors for 10 days (n = 6). Results show mean ± SD. Unpaired Student’s t test. **G, H.** Left, representative IF for CK19 (cyan), CLU (yellow), phospho-ERK1/2 (red) and DAPI (blue) (**G**) and for CK19 (red), amylase (white), Ki67 (green) and DAPI (blue) (**H**) conducted on pancreatic samples from KC mice treated with cerulein for 2 days and the indicated inhibitors for 10 days (n = 6). Scale bar, 2 mm. Right, quantification of staining plotted as mean ± SD. Unpaired Student’s t test.

Dose escalation with the FGFR2 inhibitor lirafugratinib reduced the growth of gefitinib-treated control but not *Fgfr2* knock-out T^m/+^ organoids in a dose-dependent manner, confirming the specificity of this small molecule for FGFR2 (Supplementary Fig. S8B) (44). To investigate MAPK signaling downstream of FGFR2, we induced FGFR2 up-regulation by inhibiting EGFR via erlotinib treatment for 2 days in T^m/+^ organoids and acutely inhibited FGFR2 for 1 hour with lirafugratinib. While lirafugratinib treatment alone did not affect ERK1/2 phosphorylation, sequential treatment with erlotinib followed by lirafugratinib resulted in a significant reduction in ERK1/2 phosphorylation (Fig. 7D). In line with the genetic results, FGFR2 inhibition by lirafugratinib did not affect T^m/+^ organoids growth; however, combined inhibition of FGFR2 and EGFR by lirafugratinib and erlotinib, respectively, further reduced cell growth compared to EGFR inhibition alone (Fig. 7E; Supplementary Fig. S8C). Together, these results indicated that combined inhibition of FGFR2 and EGFR reduces MAPK signaling and proliferation in pre-neoplastic organoids.

To test the cooperation between FGFR2 and EGFR in promoting pancreatic tumorigenesis, we performed an *in vivo* study. Two-month-old KC mice were treated with cerulein to expedite tumorigenesis and concomitantly received lirafugratinib and erlotinib alone or in combination (Supplementary Fig. S8D). The study was limited to 10 days, since longer term treatment led to excessive weight loss in the combination arm (Supplementary Fig. S8E). Even in this short course of treatment, we observed a significant reduction in the fraction of diseased pancreatic tissue in the mice that received both lirafugratinib and erlotinib compared to vehicle-treated mice, which was confirmed by immunolabeling for CLU, CK19 and amylase (Fig. 7F-H; Supplementary Fig. S8F). Notably, analysis of phosphorylated ERK1/2 by IF revealed a significant decrease upon erlotinib treatment alone or the combination with lirafugratinib (Fig. 7G; Supplementary Fig. S8F). However, only the combination therapy was able to significantly reduce the percentage of Ki67-positive cells (Fig. 7H). These results indicated that EGFR and FGFR2 cooperate to facilitate mutant KRAS-driven tumorigenesis by promoting proliferation and MAPK pathway activation.

## DISCUSSION

PanINs, the precursor lesions of PDA, are common in the general healthy population and rarely develop into carcinoma through less-defined mechanisms (3). An increase in gene dosage of mutant *KRAS* was implicated in driving early tumorigenesis, suggesting that neoplastic progression may be initiated when KRAS signaling exceeds a critical threshold (45). In lung cancer, activation of p53 was triggered only by enhanced oncogenic KRAS signaling, driving the selective pressure against p53 tumor suppressive function (46). Interestingly, 3D modeling of human PanINs revealed the expansion of a clonal *TP53* mutation as low-grade progressed to high-grade PanINs (2). Moreover, analysis of mouse models indicated that pancreas-specific oncogenic KRAS expression in organogenesis gave rise to a normal, functional pancreas and only sporadically promoted ADM and mPanINs formation (17,47). Furthermore, these mPanINs progressed to PDA at low frequency. Inflammatory conditions enhanced pancreatic epithelial cell plasticity and selected for subpopulations with increased mutant KRAS activity which initiated malignant invasion (48). Altogether, these findings indicated that disease progression is determined by mutant KRAS signaling intensity.

Here, we discovered a novel role for the RTK FGFR2 in enhancing mutant KRAS signaling in early pancreatic tumorigenesis. We found that FGFR2 was induced as soon as KRAS^G12D^- initiated malignant transformation of acinar cells but was neither expressed in the healthy pancreas nor after injury. Inactivation of *Fgfr2* reduced KRAS^G12D^ signaling activity measured by phosphorylation of ERK1/2 in acinar-enriched epithelial explants and the spontaneous as well as the cerulein-accelerated formation of pre-neoplastic lesions in *Kras*^G12D^ mice. Notably, the surrounding fibroblasts provided the ligands necessary for FGFR2 activation, highlighting the importance of a pre-cancer fibroinflammatory microenvironment to enhance mutant KRAS activity and initiate transformation.

RTKs present substantial overlap in signaling cascades, allowing for the augmentation of downstream signaling. The RTK EGFR is required for oncogenic KRAS-driven malignant transformation of acinar cells and is currently being explored as a potential cancer interception target (4,5,49). Importantly, here we found that FGFR2 was up-regulated and promoted the growth of pre-cancerous organoids upon EGFR inhibition. Similar results were previously described in non-small cell lung cancer (50). Reciprocally, EGFR was shown to mediate resistance to FGFR2 inhibition in FGFR2 fusion-positive cholangiocarcinoma (51).

Using a mouse model of early pancreatic tumorigenesis, we showed that only combined inhibition of EGFR and FGFR2 was able to reduce mutant KRAS-driven tumorigenesis and proliferation of pancreatic cells.

With the increasing number of FGFR2 inhibitors entering the clinic, this study lays the foundation to explore the potential use of these compounds in combination with EGFR inhibitors for PDA interception. However, further work is required to determine the target population with high risk of developing pancreatic cancer that should receive these therapies and potential adaptative escape mechanisms associated with the redundancy of RTKs signaling.

## METHODS

### Animals

*Kras^LSLG12D^, Trp53^LSLR172H^, Pdx1-Cre, Rosa26^LSLYFP^* and *Fgfr2^flox^* strains in C57Bl/6 background were interbred to obtain *Pdx1-Cre* (C); *Pdx1-Cre; Fgfr2^flox^*(C *Fgfr2^f^*); *Pdx1-Cre; Rosa26^LSLYFP^* (CY); *Kras^LSLG12D/+^; Pdx1-Cre* (KC); *Kras^LSLG12D/+^; Pdx1-Cre; Fgfr2^flox^* (KC *Fgfr2^f^*); *Kras^LSLG12D/+^; Pdx1-Cre; Rosa26^LSLYFP^* (KCY); *Kras^LSLG12D/+^; Trp53^LSLR172H/+^; Pdx1-Cre* (KPC); *Kras^LSLG12D/+^; Trp53^LSLR172H/+^; Pdx1-Cre; Fgfr2^flox^* (KPC *Fgfr2^f^*) and *Kras^LSLG12D/+^; Trp53^LSLR172H/+^; Pdx1-Cre; Rosa26^LSLYFP^*(KPCY) mice (17,18,21,39). C57Bl/6 were bred in house. All animal experiments were conducted in accordance with procedures approved by the IACUC at Cold Spring Harbor Laboratory (CSHL).

### Pre-neoplastic and neoplastic cells isolation from murine tumors

Primary KPC or KPCY tumor tissues were carefully dissected, avoiding adjacent normal pancreas or other tissue contamination. Tumors were minced and digested for 1 hr at 37°C in digestion buffer (DMEM, 5% FBS, penicillin, streptomycin, 2.5 mg/ml Collagenase D (Sigma), 0.5 mg/ml Liberase DL (Sigma), 0.2 mg/ml DNase I (Sigma)) while shaking. Single-cell suspensions were obtained by filtering through 100 µm nylon cell strainers and subsequent hypotonic lysis of red blood cells using ACK lysis buffer (Gibco). Pre-neoplastic and neoplastic KPC cells were enriched using mouse Tumor Cell Isolation Kit (Miltenyi Biotech) by magnetic cell sorting (MACS), according to manufacturer’s instructions. To isolate pre-neoplastic and neoplastic KPCY cells, cells were stained with 1 μg/ml DAPI and DAPI-negative YFP-positive cells were sorted with the FACSAria cell sorter (BD). Pre-neoplastic and neoplastic cells were prepared for subsequent experiments or seeded in growth-factor-reduced (GFR) Matrigel (BD).

### Murine Pancreatic Ductal Organoid Culture

Organoids were established as described in (25). T^m/+^ and T^m/LOH^ organoid pairs were derived from passage 1 organoids (25). Cells were seeded in GFR Matrigel (BD). When indicated, GFR Matrigel (BD) was mixed with rat tail collagen I (ThermoFisher) prepared as described in (52). Once Matrigel was solidified, pancreatic organoid medium was added. Pancreatic organoid medium (“Complete medium”) contains AdDMEM/F12, 10 mM HEPES (Invitrogen), Glutamax 1X (Invitrogen), penicillin/streptomycin 1x (Invitrogen), 500 nM A83-01 (Tocris), 50 ng/ml mEGF, 100 ng/ml mNoggin (Peprotech), 100 ng/ml hFGF10 (Peprotech), 10 nM hGastrin I (Sigma), 1.25 mM N-acetylcysteine, 10 mM Nicotinamide (Sigma), B27 supplement 1X (Invitrogen), R-spondin conditioned medium (10% final). Pancreatic organoid minimal medium (“Minimal medium”) contains AdDMEM/F12, 10 mM HEPES (Invitrogen), Glutamax 1X (Invitrogen), penicillin/streptomycin 1x (Invitrogen). For the isolation of T^m/LOH^ organoids, cells were passaged 4 times in Complete medium supplemented with 10 μM Nutlin-3a (Sigma).

For RT-qPCR assessment of TGFβ-induced transcriptional changes, T^m/+^ organoids were passaged by dissociating and reseeding into Matrigel droplets. A portion of the cells were cultured with Complete medium, while a distinct portion of passage-matched cells were cultured in Minimal medium. Cells were cultured for 7 days in Minimal medium and then treated for 24-48 hours with 5 ng/ml Recombinant Mouse TGFβ1 (R&D), unless otherwise specified, or vehicle control before being harvested for RNA extraction.

For RT-qPCR assessment of transcriptional changes upon *Gata6* knock-down, T^m/+^ organoids were cultured for 24 hours in Complete medium supplemented with 1 μg/ml doxycycline (Sigma) before being harvested for RNA extraction.

For Fig. 7A, T^m/+^ organoids were cultured for 1 day in pancreatic organoid reduced medium (no mEGF) and then treated for 48 hours with 1 μM gefitinib (Selleckchem), 1 μM erlotinib (MedChem) or DMSO control before being harvested for protein extraction.

For Fig.7D, organoids were grown for 2 days in pancreatic organoid reduced medium (no mEGF) supplemented with 1 μM erlotinib (MedChem) or vehicle. Organoids were harvested for protein extraction 1 hour after treatment with 100 nM lirafugratinib (MedChem) or vehicle.

For dissociating organoids into single cell suspensions, organoids were incubated in TrypLE Express Enzyme (ThermoFisher) for 15 min while shaking. To generate KPC-2D cell lines from tumor organoid cultures, organoids were dissociated into single cells as described above, resuspended with DMEM supplemented with 5% FBS, penicillin and streptomycin and plated on tissue culture plates. Bright Field pictures of organoids were taken with a Nikon eclipse TE2000- S microscope.

### PCR-based genotyping of *Kras, Trp53* and *Fgfr2* alleles

Genomic DNA from mouse tail and pancreas specimens was extracted with DNEasy Blood & Tissue Kit (Qiagen) following the protocol for tissue digestion. Organoids were harvested from two wells of 24 well plate and centrifuged at 1500 rpm for 5 min at 4C. Genomic DNA from freshly isolated neoplastic cells or organoids was extracted with DNEasy Blood & Tissue Kit (Qiagen) following the protocol for cultured cells.

Each PCR reaction for p53 1loxP genotyping was performed in a 20 μl mixture containing 1x AmpliTaq Gold 360 master mix (ThermoFisher), 0.5 μM each primer and 40 ng template DNA. The following primers were used: *For* AGCCTGCCTAGCTTCCTCAGG, *Rev* CTTGGAGACATAGCCACACTG. The PCR cycling conditions were 95°C for 5 minutes, followed by 40 cycles at 95°C for 30 seconds, 56°C for 30 seconds, and 72°C for 30 seconds, with a final extension step at 72°C for 5 minutes.

Each PCR reaction for genotyping of *Fgfr2* alleles was performed in a 25 μl mixture containing 1x Taq Master Mix (Dye Plus) (Vazyme), 0.4 μM each primer and 50 ng template DNA. The following primers were used: F1 ATAGGAGCAACAGGCGG, F2 TGCAAGAGGCGACCAGTCAG and F3 CATAGCACAGGCCAGGTTG. F1 and F2 produced a 142 bp and a 207 bp fragment from wild-type and *Fgfr2^flox^* alleles, respectively. F1 and F3 produced a 471 bp fragment from the *Fgfr2*^Δ^ allele. The PCR cycling conditions were 95°C for 5 minutes, followed by 35 cycles at 95°C for 30 seconds, 60°C for 15 seconds, and 72°C for 45 seconds, with a final extension step at 72°C for 5 minutes.

Each PCR reaction for genotyping of *Kras* alleles was performed in a 20 μl mixture containing Advantage GC Polymerase (Takara), 1x GC 2 PCR Buffer, 0.5 M GC-Melt, 0.5 μM dNTPs, 1.25 μM each primer and 50 ng template DNA. The following primers were used: *For* GGGTAGGTGTTGGGATAGCTG, *Rev* TCCGAATTCAGTGACTACAGATGTACAGAG. Primers produced a 285 bp and a 315 bp fragment from wild-type and *Kras^G12D^* alleles, respectively. The PCR cycling conditions were 98°C for 5 minutes, followed by 35 cycles at 98°C for 30 seconds, 58°C for 30 seconds, and 72°C for 30 seconds, with a final extension step at 72°C for 5 minutes.

PCR products were separated on a 2% agarose gel in 1x TAE buffer. Gel imaging was performed with a Syngene UV transilluminator or a ChemiDoc Imaging System (Bio-Rad).

### DNA content analysis by Propidium Iodide staining

Organoids were harvested from two wells of 24 well plate and dissociated into single cells by incubating them in TrypLE Express Enzyme (ThermoFisher) for 15 min while shaking. To analyze DNA content profile, 1×10^5^ freshly isolated neoplastic cells or organoid cell suspensions were resuspended in 1 ml of PBS and fixed by adding 2 ml of ice-cold absolute ethanol and kept at 4°C for at least 30 minutes. Cells were washed with 1 ml of 1% Bovine Serum Albumine (BSA) in PBS and stained overnight with 50 μg/ml Propidium Iodide and 250 μg/ml RNaseA at 4°C. All fluorescence-activated cell sorting (FACS) data were acquired using a LSRFortessa cell analyzer (BD) and analyzed with FlowJo software (TreeStar).

### Cell lines culture

HEK293T and KPC-2D cells were cultured in DMEM supplemented with 5% FBS, penicillin and streptomycin.

### Plasmids

Cas9-expressing T^m/+^ organoids were established by infection with LentiV_Cas9_puro vector (Addgene_108100). For knockout experiments, sgRNAs were cloned into LRG2.1_Neo (Addgene_125593). Sequences of sgRNAs were TGCATCGAAAGGCAACCTCC for *Fgfr2* KO1, GGAATACCTCCGAGCCCGG for *Fgfr2* KO2 and GAAGATGGGCGGGAGTCTTC for *Rosa26*. For knock-down experiments, shRNAs were cloned into LT3GEPIR (Addgene_111177). Sequences of shRNAs were: shGata6_1657 TGGAGTTTCATATAGAGCCCGC, shGata6_2857 TTTTCTTTTAAACAATTGGGAA, shGata6_3085 TAATGTAAACCAACCTGTGGGT, shRluc CAGGAATTATAATGCTTATCTA. FLAG-tagged mouse *Fgfr2* cDNA was cloned into LentiV_Blast (Addgene_111887) using Gibson assembly (NEB).

### Virus production and transduction

Lentivirus was produced in HEK293T cells using helper plasmids (VSVG and psPAX2) with X- tremeGENE 9 DNA Transfection Reagent (Roche). Organoids were dissociated into single cells by incubating them in TrypLE Express Enzyme (ThermoFisher) for 15 min while shaking and spin infected with the virus and 10 µg/ml polybrene (1700rpm for 45 min at room temperature). For organoid infection, the lentiviral supernatant was concentrated 10 times with Lenti-X concentrator (Takara). Media were changed at 24 hours after infection and antibiotics (2 µg/ml puromycin, 1 mg/ml G418 or 10 µg/ml blasticidin) were added at 48 hours after infection.

### *In vitro* viability/growth assay

Organoids were dissociated into single cells by incubating them in TrypLE Express Enzyme (ThermoFisher) for 15 min while shaking. Cells were counted and diluted to 10 cells/μl in a mixture of pancreatic organoid medium (90% final concentration) and GFR Matrigel (BD, 10% final concentration). 150 μl per well of this mixture (1500 cells per well) was plated in 96-well white plates (Nunc), whose wells had been previously coated with poly(2-hydroxyethyl methacrylate) (Sigma) to prevent cell adhesion to the bottom of the wells. Cell viability was measured every 24 hr, starting one day after plating, using the CellTiter-Glo assay (Promega) and SpectraMax I3 microplate reader (Molecular Devices).

For assessment of growth changes upon drug treatment, cells were diluted to 30 to 50 cells/μl in a mixture of pancreatic organoid reduced medium (no mEGF) (90% final concentration) and GFR Matrigel (BD, 10% final concentration). 1 μM lirafugratinib (MedChem) or 1 μM erlotinib (MedChem) were added 24 hours post plating using a HP D300 Digital Dispenser. Compounds were dissolved in DMSO and all treatment wells were normalized for DMSO content. Cell viability was assessed after 3 days of treatment using the CellTiter-Glo assay (Promega) and SpectraMax I3 microplate reader (Molecular Devices).

### Dose-response curves

Organoids were dissociated into single cells by incubating them in TrypLE Express Enzyme (ThermoFisher) for 15 min while shaking. 500 cells/well were plated in 384-well white plates (Nunc) in 30 μl of pancreatic organoid reduced medium (no mEGF) (90% final concentration) and GFR Matrigel (BD, 10% final concentration) and supplemented with 1 μM gefitinib (Selleckchem). Lirafugratinib was added 24 hours post plating using a HP D300 Digital Dispenser. Lirafugratinib was tested in 6 replicate wells/concentration: ranging from 1×10^-10^ M to 5×10^-5^ M. Lirafugratinib was dissolved in DMSO and all treatment wells were normalized for DMSO content. After 3 days cell viability was assessed using CellTiter-Glo as per manufacturer’s instruction (Promega) on a SpectraMax I3 (Molecular Devices) plate reader. Changes in viability were assessed relative to DMSO treated cells.

### 3D culture of KC acinar-enriched explants

3D culture of KC acinar-enriched explants in collagen was performed following the protocol described in (41). Briefly, pancreas from KC *Fgfr2^+/+^* and *Fgfr2^f/f^*mice aged 6 to 10 weeks was harvested and rinsed twice in 5 ml cold HBSS (Gibco). Tissue was minced into 1-3mm sized pieces then centrifuged for 2 min at 300 g and 4C. The buffer was aspirated and minced tissue was digested in 5 ml cold HBSS adding 100 μl 10mg/ml Collagenase P (Roche) for 15-20 minutes, shaking at 300 rpm at 37°C. During this time, mechanical dissociation by pipetting up and down was performed every 5 minutes. Collagenase P was inhibited by addition of 5 ml cold 5% FBS in HBSS. Cells were centrifuged for 2 minutes at 300 g and 4°C then washed three times with 5 mL cold 5% FBS in HBSS. Cells were passed through 500 μm strainer (Pluriselect), then through a 100 μm cell strainer (Corning), and then pelleted through 10 ml of 30% FBS in HBSS gradient. Cells were resuspended in media and incubated at 37°C for at least 4 hours prior to plating. Cells were cultured in 1x RPMI1640 supplemented with 0.1 mg/mL soybean trypsin inhibitor (ThermoFisher), 1 ug/ml dexamethasone (Sigma), 1% FBS and pen/strep. All media was sterilized through a 0.22 μm filter (VWR; Corning). Acinar-enriched clusters were processed for RNA and protein extraction or embedded in rat tail collagen I (ThermoFisher) and plated in collagen-coated 24-well plates. Culture media was added on top of solidified matrix and changed on days 1 and 3 after plating. Cells/collagen discs were digested in HBSS adding 1:50 10mg/ml Collagenase P (Roche) for ∼30 minutes, shaking at 300 rpm at 37°C. After all of the collagen was digested, cells were centrifuged for 2 minutes at 300 g and 4°C then washed with HBSS. Cells were then processed for RNA and protein extraction. Cells/collagen disks were fixed with 10% neutral buffered formalin for 15 minutes, followed with 70% ethanol overnight, and then processed for histology.

Bright Field pictures the cultures were taken with an Echo Laboratories RVL-100-G microscope.

### Western blotting

Cells were lysed with RIPA Buffer (300 mM NaCl, 5 mM EDTA, 20mM HEPES, 10% glycerol, 1% Triton X-100) supplemented with protease inhibitors (cOmplete, Mini, EDTA-free Protease Inhibitor Cocktail, Roche) and phosphatase inhibitors (PhosSTOP, Roche). Cleared lysates were electrophoresed and immunoblotted with the indicated primary antibodies: FGFR2 (CST, 23328 clone D4L2V), COFILIN (CST, 5175 clone D3F9), AMYLASE (Abcam, ab21156), phospho-p44/42 MAPK (ERK1/2) (Thr202/Tyr204) (CST, 4370 clone D13.14.4E), ERK1/2 (CST, 9102), EPCAM (CST, 93790 clone E6V8Y), β-ACTIN (CST, 4970 clone 13E5). After incubation of the membranes with appropriate HRP-conjugated secondary antibodies (Jackson ImmunoResearch Laboratories), imaging was performed using an enhanced chemiluminescence (ECL) detection kit (Cytiva) and autoradiography films (LabScientific).

### *In vivo* transplantation assay

Organoids were cultured as described above and quickly harvested on ice in AdDMEM/F12 medium supplemented with HEPES 1x (Invitrogen), Glutamax 1x (Invitrogen), and penicillin/streptomycin (Invitrogen). Organoids were dissociated to single cells with TrypLE Express Enzyme (ThermoFisher). Cells were resuspended in 50 μl of GFR Matrigel (BD) diluted 1:1 with cold PBS. Mice were anesthetized using Isoflurane and subcutaneous administration of Ketoprofen (5 mg/kg). The surgery site was disinfected with Iodine solution and 70% ethanol. An incision was made in the upper left quadrant of the abdomen. 10^5^ cells were injected in the tail region of the pancreas of wild-type C57Bl/6 mice. The incision at the peritoneal cavity was sutured with Coated Vicryl suture (Ethicon) and the skin was closed with wound clips (Reflex7, CellPoint Scientific Inc). Mice were euthanized 39 days after surgery, at which point the mice transplanted with T6^m/LOH^ organoids had reached humane endpoint. Tumors were dissected out and processed for histology.

### Cerulein treatments

All mice aged 6 to 10 weeks were food deprived for 16 hours prior to the start of the study and given eight-hourly intraperitoneal injections of 80 μg per kg of caerulein (Sigma) or saline for 2 consecutive days. Mice were weighed prior to the start of the study.

### *In vivo* therapeutic study

KC mice aged 6 to 10 weeks were randomly assigned to the four treatment groups. All mice were food deprived for 16 hours prior to the start of the study and given eight-hourly intraperitoneal injections of 80 μg per kg of caerulein (Sigma) for 2 consecutive days. Concomitantly, mice were treated via oral gavage for 10 consecutive days with 50 mg/kg erlotinib (MedChem) every 24 hours, 30 mg/kg lirafugratinib (MedChem) every 12 hours, the combination or 0.5% methylcellulose + 0.1% Tween 80 as vehicle control. Mice were weighed prior to the start of the study and every 3 days.

### Histology

All tissues were fixed with 10% neutral buffered formalin overnight. Tissues were then processed with standard tissue processing protocol, embedded in paraffin and 6 μm sections were cut and mounted on slides. Formalin fixed paraffin-embedded (FFPE) tissue sections were stained with hematoxylin and eosin (H&E) or used for immunohistochemical/immunofluorescent labeling.

Analysis of diseased pancreas was performed using QuPath software (53) for percentage area on a digital image of the tissue section acquired with Aperio scanner (Leica Biosystems) or Olympus VS200 scanner. H&E sections were evaluated for highest grade lesion present in a blinded fashion.

The human PDA and IPMN tissue microarrays used for IF and IHC were kindly provided by the Sidney Kimmel Comprehensive Cancer Center at Johns Hopkins University. The human NAT tissues were kindly provided by Mount Sinai School of Medicine biorepository core facility. Human PanIN tissue microarrays were obtained from US Biomax (BIC14011b). All tissue donations and experiments were reviewed and approved by the Institutional Review Board of Cold Spring Harbor Laboratory and the clinical institutions involved. Written informed consent was obtained prior to acquisition of tissue from all patients, or consent was waived by local institutional review boards. The studies were conducted in accordance with ethical guidelines (Declaration of Helsinki). Samples were confirmed to be PanIN, IPMN or PDA on pathologist assessment.

### Immunohistochemistry (IHC), immunofluorescent labeling (IF), RNA *in situ* hybridization (RNA ISH) and RNA ISH combined with IF

FFPE sections were deparaffinized and rehydrated. For antigen retrieval, slides were incubated with 10 mM Citrate buffer (pH 6.0) in a pressure cooker for 6 minutes. To perform IHC, endogenous peroxidase activity was quenched in 3% H_2_O_2_ for 20 min. Tissues were blocked in 2.5% Normal Horse Serum blocking solution (Vector Laboratories) for IHC or 5% BSA (Sigma) in TBST buffer for IF and subjected to staining with the following antibodies overnight at 4C: p53 (Leica, P53-CM5P-L 1:100), GFP/YFP (Abcam, ab6673 1:100), AMYLASE (Sigma, A8273 1:250), Dolichos Biflorus Agglutinin (DBA) Fluorescein (VectorLabs, FL-1031 1:200), FGFR2 (Abcam, ab58201 1:1000), CK19 Alexa Fluor 647 (Abcam, ab205446 clone EPR1579Y 1:500), CK19 (Sigma, MABT913 1:500), GATA6 (R&D, AF1700 1:500), S100A2 (Abcam, ab109494 1:250), FGF7 (Origene, TA321423 1:100), FGF10 (Sigma, ABN44 1:100), phospho-p44/42 MAPK (ERK1/2) (Thr202/Tyr204) (CST, 4370 clone D13.14.4E 1:250), Ki67 (ThermoFisher, 14-5698-82 clone SolA15 1:500), Ki67 Alexa Fluor 488 (Abcam, ab281847 clone SP6 1:500), clusterin (ThermoFisher, PA5-46931 1:100).

ImmPRESS Alkaline Phosphatase (AP) and ImmPRESS Horseradish Peroxidase (HRP) IgG polymers (Vector Laboratories) were used as secondary antibodies for IHC. ImmPACT DAB peroxidase substrate and Vector Blue Alkaline Phosphatase substrate (Vector Laboratories) were used as substrates. Hematoxylin (Vector Laboratories) was used as counterstain. Cover slides were mounted with Cytoseal 60. Bright field images were obtained using a Zeiss Axio Imager.A2 microscope.

Secondary Alexa Flour antibodies for IF were obtained from ThermoFisher. Nuclei were stained with 1 μg/ml DAPI for 1 hr. Slides were mounted with ProLong Gold Antifade Mountant (ThermoFisher). Z-stack images were obtained using a Leica SP8 confocal laser microscope at 40x magnification. Large mosaic stitched images were obtained using a Zeiss Observer inverted fluorescence microscope. Tissue sections were imaged at 20x with an Olympus VS200 scanner.

RNA ISH combined with IF was performed according to manufacturer’s instructions (RNAscope Multiplex Fluorescent Reagent Kit v2 323100; ACD) using probes specific for *Fgfr1* (443491-C2; ACD), *Vim* (457961-C4; ACD) and *Cdkn2a/*ARF (503811-C1; ACD). Once completed, the samples were washed in PBS then blocked for 1 hour with 20% donkey serum at room temperature. Primary antibodies were incubated overnight at 4°C. Secondary Alexa Flour antibodies (1:500 in blocking buffer) were incubated for 1 hour at room temperature, and samples were washed 3 times in PBS. Slides were counterstained with DAPI and mounted with ProLong Gold Antifade Mountant (ThermoFisher). Z-stack images were obtained using a Zeiss LSM780 confocal laser scanning microscope at 20x magnification.

Quantification of staining was performed with QuPath software (53).

### RNA Extraction

RNA was extracted using the TRIzol Plus RNA Purification Kit (ThermoFisher) per manufacturer’s instructions. RNA samples were treated on column with PureLink DNase (ThermoFisher).

### Expression analysis by RT-qPCR

Total RNA was isolated as described above. Complementary DNA (cDNA) was produced using the TaqMan reverse transcription reagents (ThermoFisher). 10 ng of cDNA were used for RT-qPCR reactions with TaqMan Universal Master Mix II, no UNG (ThermoFisher) and the following TaqMan probes: *Fgfr2* Mm01269930_m1*, Gata6* Mm00802636_m1, *Fgfr1* Mm00438930_m1, *Snai1* Mm00441533-g1, *Fn1* Mm01256744_m1 and *Hprt* Mm00446968_m1 (ThermoFisher).

Z-score values were derived from 2^-ΔCt values.

### RNA sequencing

The quality of purified RNA samples was determined using a Bioanalyzer 2100 (Agilent) with an RNA 6000 Nano Kit. RNAs with RNA Integrity Number (RIN) values greater than 8.5 were used to generate sequencing libraries. Libraries were generated from 1 μg of total RNA using a TruSeq Stranded Total RNA Library Prep Human/Mouse/Rat (48 Samples) (Illumina) or KAPA mRNA HyperPrep Kit (Roche) per manufacturer’s instructions. Libraries were quality checked using a Bioanalyzer 2100 (Agilent) with a High Sensitivity DNA Kit and quantified using PicoGreen (ThermoFisher). Equimolar amounts of libraries were pooled and subjected to single- or paired-end, 75 bp sequencing at the Cold Spring Harbor DNA Sequencing Next Generation Shared Resource using an Illumina NextSeq 500 or 550 instrument.

### RNA-sequencing Data Analysis

RNA-seq reads quality was first quantified using FastQC (http://www.bioinformatics.babraham.ac.uk/projects/fastqc/). Reads were then trimmed using Trim Galore (54). Reads were mapped to transcript annotation GENCODE M16 (organoids RNA-seq) and GENCODE M35 (KC acinar-enriched explants RNA-seq) (55) using Spliced Transcripts Alignment to a Reference (STAR) (56). RSEM (57) was used to extract counts per gene and transcripts per million (TPM). RNA-seq tracks were generated from aligned reads using deepTools2 (58) and visualized using UCSC Genome Browser (59).

Differential gene expression analysis was performed using Bioconductor package DESeq2 (60). A pre-filtering step was performed to remove genes that have reads in less than two samples. T^m/+^ and T^m/LOH^ organoids were considered paired datasets. A pre-filtering step was performed to remove genes that have reads in less than two samples. At this step, all genes not classified as protein coding according to BioMart were discarded (61). Only genes with an adjusted p-value less than 0.05 were retained as significantly differentially expressed genes. A principal component analysis (PCA) was performed using the 500 most variable protein coding genes after variance stabilizing transformation (VST) using DESeq2 software. The graphical representation of the two most important components was created using CRAN package ggplot2 (62).

Default parameters were used to perform GSEA Preranked analysis using the GSEA software (63,64). Hallmark and curated gene sets were downloaded from the MSigDB database, which include pathways from the Kyoto Encyclopedia of Genes and Genomes (KEGG) (65,66).

### *Trp53* SNP Calling

SNP calling was based on the GATK RNA-seq short variant discovery pipeline (67). More specifically, the reads were grouped per sample and duplicated reads marked using Picard MarkDuplicates (http://broadinstitute.github.io/picard). Then, the GATK toolkit (68) was used, in this specific order, to span splicing events and to reassign map quality (SplitNCigarReads program), to realign indels (RealignerTargetCreator and IndelRealigner programs), to recalibrate the base quality scores (BaseRecalibrator program) and to call and filter variants (HaplotypeCaller and VariantFiltration programs) across all samples. Read count and quality were extracted, for variants of interest, using mpileup option from SAMtools (69). Using a home-made script, reads with mapping quality filter (MAPQ value) < 5 and sequencing quality score (phred value) < 20 were removed before counting the number of reads (coverage) for each allele present at the position of interest.

### Single cell RNA-sequencing Data Analysis

The scRNA-Seq plot was generated with ggplot2 (70). A cell was considered as expressing the gene when the transcription count was higher than zero. The cell clusters corresponded to the one defined in the associated paper.

### Other bioinformatic and statistical analyses

GraphPad Prism 10 was used for graphical representation of data. Heatmaps were generated using GraphPad Prism 10 or RStudio and Bioconductor package ComplexHeatmap (71,72). Statistical analysis was performed using the tools within Prism 10 indicated in the figure legends. Asterisks denote P-value as follows: *, P ≤ 0.05, **, P ≤ 0.01, ***, P ≤ 0.001, ****, P ≤ 0.0001, ns, not significant. Figures were prepared using Illustrator (Adobe).

## Data availability

RNA-seq data generated in this study have been deposited in the Gene Expression Omnibus (GEO) database under the accession number GSE278148 (KC acinar-enriched explants RNA-seq) and GSE278203 (organoids RNA-seq).

Publicly available data reanalyzed for this study are available in the GEO database under the accession code GSE195914 (pre-neoplastic and neoplastic KPC cells scRNA-seq) (31), GSE93326 (human PDAs LCM RNA-seq) (73), GSE71729 (human PDAs microarray data) (74), GSE250519 (CUT&RUN) (31), in the EGA under accession code EGAS00001002543 (human PDAs LCM RNA-seq) (35). Expression and mutational data for PDA studies were obtained from the cBioPortal (75,76) (paad_tcga (77)).

## ACKNOWLEDGEMENTS

We thank Dr. Klingbeil, Dr. Caligiuri and Dr. Shakiba for critical reading of this manuscript. This work was performed with assistance from the Cold Spring Harbor Laboratory shared resources, which are supported by the National Institutes of Health (Cancer Center Support Grant 5P30CA045508: Bioinformatics, DNA Sequencing, Flow Cytometry, Microscopy, Animal, and Animal and Tissue Imaging Shared Resources). This work was performed with assistance from the US National Institutes of Health Grant S10OD028632-01.

The authors are supported by National Institutes of Health (Cancer Center Support Grant 5P30CA045508) and the Lustgarten Foundation, where D.A. Tuveson is a distinguished scholar and Director of the Lustgarten Foundation–designated Laboratory of Pancreatic Cancer Research. D.A. Tuveson is also supported by the Thompson Foundation, the Pershing Square Foundation, the Cold Spring Harbor Laboratory and Northwell Health Affiliation, the Northwell Health Tissue Donation Program, the Cold Spring Harbor Laboratory Association, and the National Institutes of Health (5P30CA45508, U01CA210240, R01CA229699, U01CA224013, 1R01CA188134, and 1R01CA190092). This work was also supported by a gift from the Simons Foundation (552716 to D.A. Tuveson). C. Tonelli was a fellow of the American-Italian Cancer Foundation. Y. Park is supported by the National Cancer Institute (R50CA211506).

---

**Supplementary Fig. S1**

**A.** PCR-based genotyping for assessing the *Trp53* status in neoplastic-enriched cells from KPC tumors after isolation (p0) and upon passaging as organoids in medium depleted of different components (p1 to p3). **B.** PCR-based genotyping for assessing the *Trp53* status in neoplastic-enriched cells from a KPC tumor after isolation (p0) and upon passaging as organoids in matrigel supplemented with different amounts of collagen (p1 to p4). **C.** PCR-based genotyping for assessing the *Trp53* status in neoplastic-enriched cells from KPC tumors after isolation (p0) and upon passaging as organoids in medium with or without Nutlin (p1 to p4). The DNA profile of the cells was evaluated at every passage by Propidium Iodide staining. **D.** PCR-based genotyping for assessing the *Trp53* status in FACS-sorted YFP-positive cells from a KPCY tumor after isolation (p0) and upon passaging as organoids in medium with or without Nutlin (p1 to p4). **E.** Pictures of T^m/+^ at 0, 24 and 48 hours after treatment with Nutlin.

**Supplementary Fig. S2**

**A.** DNA profile of T^m/+^ (in pink) and T^m/LOH^ organoids (in blue) as determined by Propidium Iodide staining. **B.** T^m/+^ and T^m/LOH^ organoids proliferate at similar rate as measured by CellTiter-Glo luminescent cell viability assay. **C.** Percentage of wild-type and mutant *Trp53* reads as determined by RNA-seq in T^m/+^ (pink bars) and T^m/LOH^ organoids (blue bars). **D.** *Prdm1, Foxa1, Gata5, Twist1, Glul* and *Soat1* expression in T^m/+^ (pink dots) and T^m/LOH^ organoids (blue dots) as determined by RNA-seq. Results show mean ± SD. q-value as calculated by DESeq2 is indicated. **E.** UCSC Genome Browser display of RNA-seq reads in T^m/+^ and T^m/LOH^ organoids at the *Fgfr2* locus. T^m/+^ organoids tracks are colored in pink and T^m/LOH^ organoids tracks in blue.

**Supplementary Fig. S3**

**A, B.** Left, representative IF for YFP (green), FGFR2 (pink), FGF7 (**A**) or FGF10 (**B**) (yellow) and DAPI (blue) conducted on pancreata from CY mice (n = 4), mPanINs from KCY mice (n = 6) and tumors from KPCY mice (n = 6). Right, quantification of staining plotted as mean ± SD. Unpaired Student’s t test. **C.** *FGFR2* expression in Classical and Basal-like PDA samples (35,73,74). Results show mean ± SD. Unpaired Student’s t test. **D.** *Gata6* (dark grey) and *Fgfr2* (light grey) expression as determined by RT-qPCR in T6^m/+^ and T34^m/+^ organoids expressing inducible hairpins against *Gata6* or a control hairpin against Renilla luciferase (n = 3). Results show mean ± SD. Unpaired Student’s t test. **E.** UCSC Genome Browser display of H3K4me3 and GATA6 CUT&RUN peaks in HPAF-II cells at the *FGFR2* gene (31).

**Supplementary Fig. S4**

**A.** Time-course of *Fgfr2* and *Fgfr1* expression as determined by RT-qPCR in T6^m/+^ and T19^m/+^ organoids after growth in Minimal medium supplemented with TGFβ or vehicle (n = 3). Results show mean ± SD. Unpaired Student’s t test. **B.** Single color images of RNA ISH for *Fgfr1* (magenta), *Vim* (cyan) and IF for CK19 (white), FGFR2 (green), p53 (orange) and DAPI (blue) conducted on tumors from KPC mice. Scale bar, 50 μm. **E.** Single color images of RNA ISH for *Cdkn2a/*ARF (cyan), *Vim* (yellow) and IF for YFP (white), CK19 (red), FGFR2 (green), p53 (orange) and DAPI (blue) conducted on tumors from KPCY mice. Scale bar, 20 μm.

**Supplementary Fig. S5**

**A.** PCR-based analysis of the genomic DNA extracted from the tail and the pancreas of C *Fgfr2^+/+^* and *Fgfr2^f/f^* mice for assessing the *Fgfr2* alleles (*Fgfr2^wt^*and *Fgfr2^flox^*) and recombination of the *Fgfr2* locus (*Fgfr2*^Δ^). **B.** PCR-based analysis of T^m/+^ organoids derived from KPC *Fgfr2^+/+^, Fgfr2^f/+^* and *Fgfr2^f/f^*tumors for assessing the recombination of the *Fgfr2* locus. FGFR2 protein expression as determined by Western blotting. Loading control, COFILIN.

**Supplementary Fig. S6**

**A.** Left, representative H&E staining conducted on pancreatic samples from age-matched C and KC mice at different time points after saline or cerulein treatment (day −1, day 0). Scale bar, 50 μm. Right, quantification of the fraction of normal pancreas (n = 5-8). Results show mean ± SD. Unpaired Student’s t test. **B.** Left, representative IF for CK19 (red), amylase (white) and DAPI (blue) conducted on pancreatic samples from age-matched C and KC mice at different time points after saline or cerulein treatment (day −1, day 0). Scale bars are 500 μm for main images and 50 μm for insets. Right, quantification of staining plotted as mean ± SD. Unpaired Student’s t test. **C.** Representative H&E and IHC for FGFR2 conducted on a human normal adjacent tissue sample. Scale bars are 200 μm for main images and 50 μm for insets. **D.** PCR-based analysis of the genomic DNA extracted from the tail and the pancreas of KC *Fgfr2^+/+^*and *Fgfr2^f/f^* mice for assessing the *Fgfr2* alleles (*Fgfr2^wt^* and *Fgfr2^flox^*) and recombination of the *Fgfr2* locus (*Fgfr2*^Δ^). **E.** Protein expression analysis in acinar clusters and associated terminal ductal epithelium from KC *Fgfr2^+/+^* and *Fgfr2^f/f^* mice at different time points after culture in collagen, as determined by Western blotting. Loading control, COFILIN. **F.** Heatmap of z-score values of the indicated acinar and ductal genes expression as determined by RNA-seq in acinar-enriched explants from KC *Fgfr2^+/+^*and *Fgfr2^f/f^* mice at different time points after culture in collagen. **G.** GSEA signatures ‘HALLMARK E2F TARGETS’ and ‘HALLMARK MYC TARGETS V1’ are increased in acinar-enriched explants from KC *Fgfr2^+/+^* compared to *Fgfr2^f/f^* mice at day 5 of culture in collagen. NES, normalized enrichment score; FDR, false discovery rate.

**Supplementary Fig. S7**

**A, B.** Left, representative IF for CK19 (red), amylase (white) and DAPI (blue) conducted on pancreatic samples from 7- to 11-months-old KC *Fgfr2^+/+^* and *Fgfr2^f/f^* mice (n = 7) (**A**) or from KC *Fgfr2^+/+^* and *Fgfr2^f/f^* mice harvested 30 days after cerulein treatment (day −1, day 0) (n = 8) (**B**). Scale bars are 500 μm for main images and 50 μm for insets. Right, quantification of staining plotted as mean ± SD. Unpaired Student’s t test. **C.** PCR-based genotyping of T^m/+^ and T^m/LOH^ organoids derived from KPC *Fgfr2^+/+^* and *Fgfr2^f/f^* tumors for assessing the *Kras, Trp53* and *Fgfr2* status. **D, E.** Left, representative IF for CK19 (red), phospho-ERK1/2 (green) and DAPI (blue) conducted on pancreatic samples from 7- to 11-months-old KC *Fgfr2^+/+^* and *Fgfr2^f/f^*mice (n = 6- 7) (**D**) or from KC *Fgfr2^+/+^* and *Fgfr2^f/f^*mice harvested 30 days after cerulein treatment (day −1, day 0) (n = 7-8) (**E**). Scale bars are 500 μm for main images and 50 μm for insets. Right, quantification of staining plotted as mean ± SD. Unpaired Student’s t test. **F, G.** Left, representative IF for CK19 (red), Ki67 (green) and DAPI (blue) conducted on pancreatic samples from 7- to 11-months-old KC *Fgfr2^+/+^* and *Fgfr2^f/f^* mice (n = 5-7) (**F**) or from KC *Fgfr2^+/+^* and *Fgfr2^f/f^*mice harvested 30 days after cerulein treatment (day −1, day 0) (n = 7-8) (**G**). Scale bars are 500 μm for main images and 50 μm for insets. Right, quantification of staining plotted as mean ± SD. Unpaired Student’s t test.

**Supplementary Fig. S8**

**A.** Relative growth (day 4 / day 1) of T^m/+^ organoids knocked-out for *Fgfr2* or control *Rosa26* (n = 5). Results show mean ± SD. Unpaired Student’s t test. **B.** Dose-response curves for lirafugratinib in T^m/+^ organoids knocked-out for *Fgfr2* or control *Rosa26* (n = 6). **C.** Fold change of growth inhibition of T^m/+^ organoids upon treatment with vehicle, lirafugratinib, erlotinib or the combination. **D.** Schematic representation of the study design. **E.** Quantification of pancreas-to-body weight percentages normalized to day −1 in KC mice treated with cerulein for 2 days and the indicated inhibitors for 10 days (n = 6). Results show mean ± SD. **F.** Quantification of staining from Fig. 7G plotted as mean ± SD. Unpaired Student’s t test.

